# Lamin C regulates genome organization after mitosis

**DOI:** 10.1101/2020.07.28.213884

**Authors:** X Wong, VE Hoskins, JC Harr, M Gordon, KL Reddy

## Abstract

The dynamic 3D organization of the genome is central to the regulation of gene expression and developmental progression, with its disruption being implicated in various diseases. The nuclear lamina, a proteinaceous meshwork underlying the nuclear envelope (NE), provides both structural and regulatory influences on genome organization through the tethering of large inactive genomic regions, called Lamina Associated Domains (LADs), to the nuclear periphery. Evidence suggests that the A type lamins, lamins A and C, are the predominant lamins involved in the peripheral association of LADs, with these two isotypes forming distinct networks and potentially involved in different cellular processes. Here we tested whether lamins A and C have distinct roles in genome organization by examining chromosome architecture in cells in which lamin C or lamin A are specifically down-regulated. We find that lamin C (not lamin A) is required for the 3D organization of LADs and overall chromosome organization in the cell nucleus. Striking differences in the localization of lamin A and lamin C are present as cells exit mitosis that persist through early G1. Whereas lamin A associates with the nascent NE during telophase, lamin C remains in the interior surrounding nucleoplasmic LAD clusters. Lamin C association with the NE is delayed until several hours into G1 and correlates temporally and spatially with the post-mitotic NE association of LADs. Post-mitotic LAD association with the NE, and consequently global 3D genome organization, is perturbed only in cells depleted of lamin C, and not in cells depleted of lamin A. We conclude that lamin C regulates LAD dynamics after mitosis and is a key regulator of genome organization in mammalian cells. These findings reveal an unexpectedly central role for lamin C in genome organization, including both inter-chromosomal LAD-LAD segregation and LAD scaffolding at the NE.

## Introduction

Lamins encoded by *LMNA*, *LMNB1* and *LMNB2* form networks of nuclear intermediate filaments as major components of the nucleoskeleton. Lamin filaments interact with key partners, including most nuclear membrane proteins, to form nuclear lamina networks that determine nuclear mechanics, modulate signaling and dynamically organize the genome^1–6^. Lamina networks interact with large regions of transcriptionally-silent heterochromatin in each cell type and customize the 3D configuration of individual chromosomes with respect to the nuclear envelope (NE). These silent heterochromatin regions, identified operationally as Lamina Associated Domains (LADs), correspond to the ‘B’ compartment identified via HiC and related chromatin mapping strategies^7–9^. Chromatin association with the lamina, and its opposite (dynamic release as active ‘non-LAD’ or A-compartment chromatin) are particularly important for developmentally-regulated genes needed to create or maintain cell-specific identity^8,10–12^. ‘Silent’ histone modifications, including H3 lysine 9 methylation (H3K9me2/3) and H3 lysine 27 trimethylation (H3K27me3), are key components of LAD organization^6,13–16^. We previously found that A-type lamins, encoded by *LMNA*, establish and/or maintain interphase LAD configuration^6,17^. In recent years there have been further advances in understanding how LADs, chromatin looping, epigenetic modifications and liquid-liquid phase transitions of chromatin during interphase are interrelated^2,3,5–7,13–16,18–23^

In contrast, the mechanisms by which nuclear 3D structural information is faithfully transmitted and re-established through cell division are of significance and remain largely unknown. During entry into mitosis, interphase spatial genome and NE organization are lost as chromosomes condense and the NE and nuclear lamina networks disassemble, yet global chromosome and 3D-genome organization is re-established in the next interphase^8,9,16,24–30^. As cells enter anaphase, both super-resolution imaging and single cell Hi-C (a genome-wide method to detect chromatin contacts) show that LAD regions of chromosomes begin to self-aggregate as globular ‘compartments’ prior to nuclear lamina formation^9^. These LAD agglomerations slowly make their way to the nascent NE during early G1 and ultimately ‘spread’ across the lamina as they approach and interact with the nuclear periphery^9^.

One key protein in interphase LAD organization and NE function are the A-type lamins. Alternative mRNA splicing of *LMNA* produces two main somatic isoforms, lamin A and lamin C, the first 566 or 568 residues of which are identical in human and mice, respectively^31^. Lamin C has six unique C-terminal residues, whereas lamin A has an extended tail domain that undergoes four post-translational modification steps to achieve its final mature length of 646 or 647 residues, in humans and mice respectively^32^. Until recently lamin A and lamin C were assumed to be functionally redundant. Super-resolution microscopy show lamins A, C, B1 and B2 form separate but structurally inter-dependent filament networks such that removing any one (e.g. C) lamin affects the distribution and geography of the other three^33–35^. In addition to forming separate networks, several lines of evidence suggest that these isotypes might have unique roles in cellular function. For instance, lamin A confers mechanical stiffness that determines when different white blood cell lineages exit the bone marrow. In contrast, in CNS neurons lamin C is the predominant A-type lamin isoform expressed due to the selective down-regulation of the pre-lamin A transcript by miR-9^36–38^. Studies in mice that express only one A-type lamin (either mature lamin A or lamin C) also highlight potential functional differences. Mice expressing mature lamin A (no lamin C) have few if any overt phenotypes with the exception of misshapen nuclei, whereas mice that express only lamin C (no lamin A) have longer lifespans, are mildly obese and are predisposed to cancer^39–41^.

Intriguingly, both A-type lamins and the Lamin B Receptor (LBR; a nuclear membrane protein) are essential molecular tethers for heterochromatin at the NE^42^. LBR is especially important in early development (when lamin A/C expression is low), while A-type lamins are prominent later in development, perhaps explaining why lamins are thought to be dispensable for proliferation and differentiation of mouse embryonic stem cells (mESC)^43–45^. It remains unclear if lamins are necessary for robust LAD organization in mESC^43,44^. Previously we showed in fibroblasts, which are more terminally differentiated and where LBR is only minimally expressed, interphase LAD organization is disrupted by depleting both A-type lamins (A and C), but not lamin A alone^6^. Taken together, these data suggests lamin C might be required to tether LADs at the NE in cells lacking LBR. Therefore, in this study we examine the role of lamin C in LAD organization by examining LAD and chromosome configuration in cells specifically depleted in either lamin C or lamin A. Our data strongly support the hypothesis that lamin C is uniquely required for large scale chromosome organization. Our results provide insight into the mechanisms of 3D genome organization during interphase and its dynamic re-establishment after mitosis.

## Results

### LAD proximity to the nuclear envelope is maintained by lamin C

To test our hypothesis, we developed short hairpin RNAs (shRNA) that specifically down-regulate lamin A (shA) or lamin C (shC). We also used shRNAs that down-regulate both lamins A and C (shAC), or lamin B1 (shB1)^6^. To characterize the efficacy of lamin depletion and global effects on mouse embryonic fibroblasts (MEFs), we assayed for the presence of lamin isotypes after four days and monitored growth of cells during the same time-frame (Fig. S1). Each shRNA specifically targeted its own lamin, as shown by western analysis (Fig. S1A). shC, shB1 and shlacZ (control) all grew at the same rate, while shA and shAC displayed reduced growth rates, suggesting a unique role for lamin A in cell growth and cycling (Fig S1B). To try to assess if these isotypes had effects on LAD organization, particularly at specific regions, we performed a global DamID-seq analysis of LAD positioning (using Dam-lamin B1) in shRNA treated MEFs. Surprisingly, these analyses revealed no significant differences in cells depleted of both lamin A and lamin C compared to wild-type cells (shAC; Fig. S2). While this was initially surprising, especially given our earlier findings that lamin A/C is required to organize regions to the lamina, we realize that an inherent limitation of DamID and related techniques is that data are aggregated from millions of cells, potentially obscuring true differences that would be detectable at the level of individual cells^6,9,46^. Detailed analysis of LAD boundaries (region of transition from NE-associated to NE non-associated), which are normally are quite sharp, showed that LAD boundaries also remained intact under all four conditions (control, shAC, shA, shB1, or shC); genome-wide comparisons between log_2_ ratios of DamID-seq signals were virtually indistinguishable (Fig. S2A,B and S3), and genome-wide bioinformatically defined LADs for all three down-regulated conditions showed preservation of WT LADs (>90% by base coverage) (Fig. S2C).

To observe LAD organization in single cells, we used 3D-immunoFISH to examine individual chromosome organization in shRNA down-regulated MEFs. We highlighted the NE in green using antibodies to lamin B1 (or lamin A/C in the case of shB1), and the LAD and non-LAD regions of chromosome 11 were ‘painted’ red and cyan, respectively, using oligonucleotide-based Chromosome Conformation Paints (CCP)^8,9^. As previously shown, in control nuclei, each chromosome occupied its own territory (Fig. 1), with LADs clustered together near the NE and non-LADs extending into the nucleoplasm^8,9^. This organization was grossly disrupted in cells depleted of both lamin A and lamin C (shAC); LADs failed to co-associate, with many LADs mis-localized to the nucleoplasm, and non-LADs dispersed to occupy a territory considerably larger than controls (Fig. 1). This finding is consistent with another study that showed chromosome territory expansion upon removal of lamin A/C and our finding that lamin A/C is required for de novo LAD tethering^6,17^.

**Figure 1:**
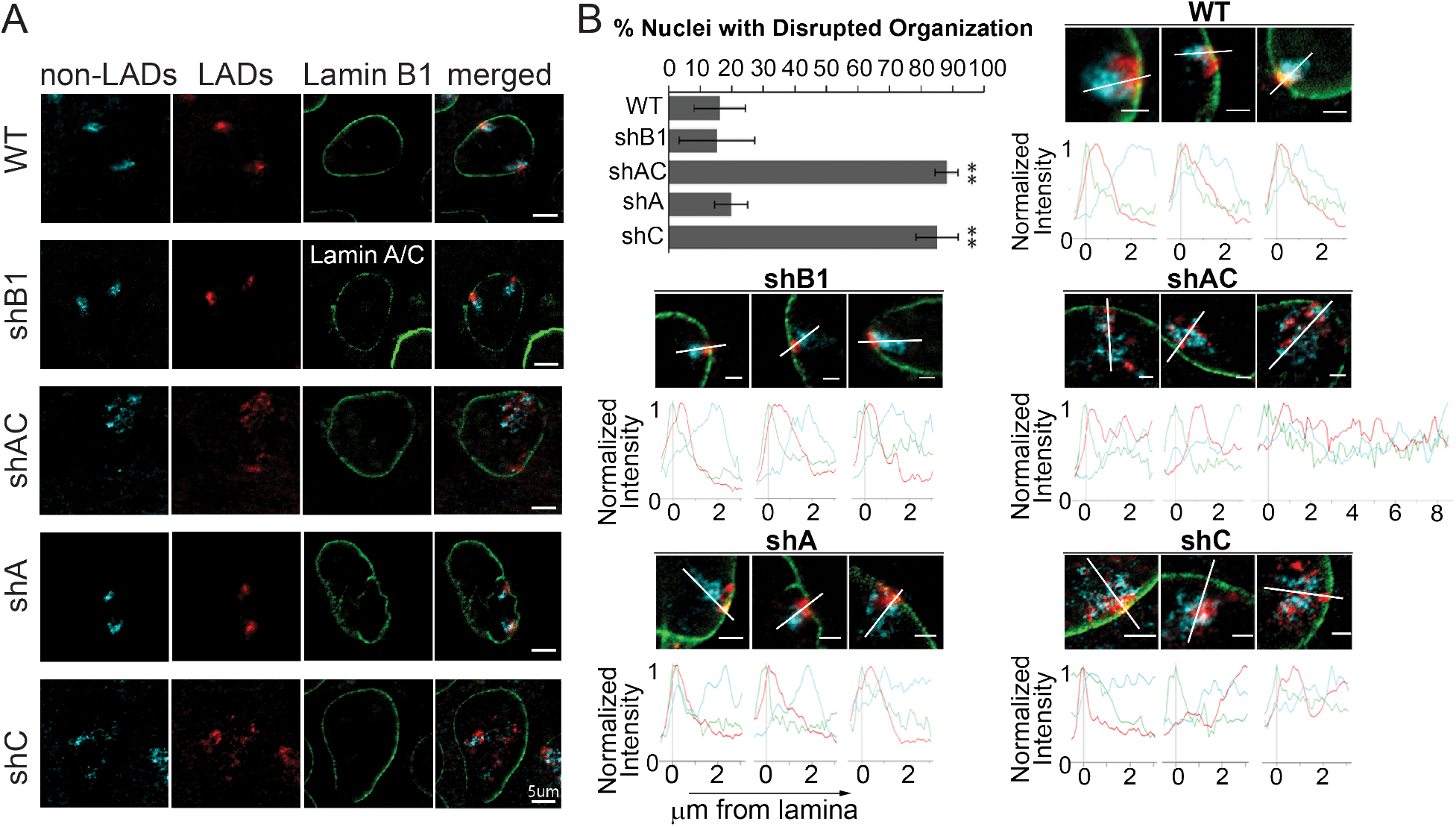
Lamin C is important for chromosomal sub-domain organization. (A) Representative images showing the organization of chromosome 11 with nonLADs in cyan, LADs in red and lamin B1 (or lamin A/C for shB1 treated cells) in green. (B) Quantification of nuclei with disrupted organization for each knockdown condition. Error bars represent 1 standard deviation. ∗ ∗ indicates t-test p value <0.001 (n>200 nuclei per condition). Representative images of chromosome conformation paints to chromosome non-LADs (cyan) LADs (red) Lamin B1 (green; lamin A/C for shB1 treated cells) in primary MEFs. Images were chosen to represent the spectrum of phenotypes for each knockdown. Normalized fluorescence intensity histogram plots: from nuclear lamina (0*μ*m) to 3*μ*m into the nucleus, were plotted for all chromosome 11 territories (except territory 3 of shAC, which was expanded to display the extent of LAD disruption for that nucleus). The line each plot travels through is represented by a white line. Scale bar = 2*μ*m.

We next asked what the role and contribution of individual lamin isotypes were to this organization. We observed no gross perturbations of genome organization after loss of Lamin B1. Furthermore, loss of lamin A alone had no discernible effect on LAD organization, even though loss of this isotype displayed delayed growth. Strikingly, loss of lamin C was sufficient to fully recapitulate the gross disruption of chromosome 11 organization seen with loss of both isotypes (Fig. 1A, shAC vs. shC). To better visualize and describe the observed perturbations, we plotted signal intensities from individual nuclei across lines drawn through the medial plane of each chromosome territory, as previously described^9^ (Fig. 1B, S4-8). This analysis confirmed the normal positioning of LADs and non-LADs in control nuclei (Fig. 1A, B, S4), lamin B-depleted nuclei (Fig. 1A,B, S5) and lamin A-depleted nuclei (Fig 1A,B and S7), and confirmed the broadening of the distribution of the LAD signal intensity and the overall dispersion of LADs away from the NE in lamin C-depleted cells (Fig. 1A, B, S8), however, because of the severity of the phenotype and the inability to discriminate between individual deranged chromosomes, we could not reliably average these data. Therefore, to accurately account for all disrupted cells, we counted the percentage of cells with disrupted genome organization in each population (Fig. 1B). Taking a conservative approach, we scored nuclei in which the majority of the LAD signal localized near the medial plane. Nuclei were considered disrupted if one (or both) chromosomes had LADs that were either dispersed (not aggregated or loss of LAD:LAD cohesion) or gross loss of NE-association. These were scored by two independent observers (blinded). By this metric, genome organization was disrupted in 16% of wild-type control nuclei (Fig. 1B). Cells depleted of lamin A showed a similar baseline (18%; Fig. 1B). We note that these cells are unsynchronized, so any alterations in LAD organization due to cell cycle stage are encompassed in this baseline data. In contrast, genome organization was disrupted in 85% of lamin C-depleted cells (p<0.001) and 88% of shAC cells (p<0.001; Fig. 1B and S6). One possibility, that lamin C was uniquely required for cell cycle progression, is unlikely since the doubling times for shC matched control cells were the same (Fig. S1, shLacZ vs. shC). Overall, these results suggest that lamin C is required for LAD self-association (LAD:LAD cohesion), LAD retention near the NE and overall compaction of the chromosome territory (including non-LAD regions).

### Lamin C (not lamin A or lamin B1) is required to maintain LAD association with the NE

To independently evaluate the role of lamin C, we used cells bearing a single ‘TCIS’ LAD. TCIS (Tagged Chromosomal Insertion Site) is comprised of 256 tandem copies of the *lacO* sequence (allowing visualization upon expression of EGFP-LacI) and a modified RMCE (recombination-mediated cassette exchange) ‘cassette’, allowing for insertion of ectopic sequences. We previously showed, using this system, that a single segment of DNA (lamina associated sequence or LAS) can redirect the TCIS locus to the nuclear lamina. To test if lamin C is required for localization of a de novo LAD, we used two independent MEF clones bearing one of these TCIS-LAS, specifically, the Ikzf1 (Ikaros zinc finger protein) D6 Lamin Associated Segment, as previously described^6^. When this LAS is introduced into the clonal TCIS MEF lines (clone Y or clone 12, Fig. 2), the TCIS-LAS locus was NE-associated in 75-80% of nuclei, compared to 40% of nuclei with a TCIS locus alone (no LAS), when visualized by 3D-immunoFISH (Fig. 2A) and quantified by co-localization with lamin B1 (or lamin A/C for shB1) (Fig. 2).

**Figure 2:**
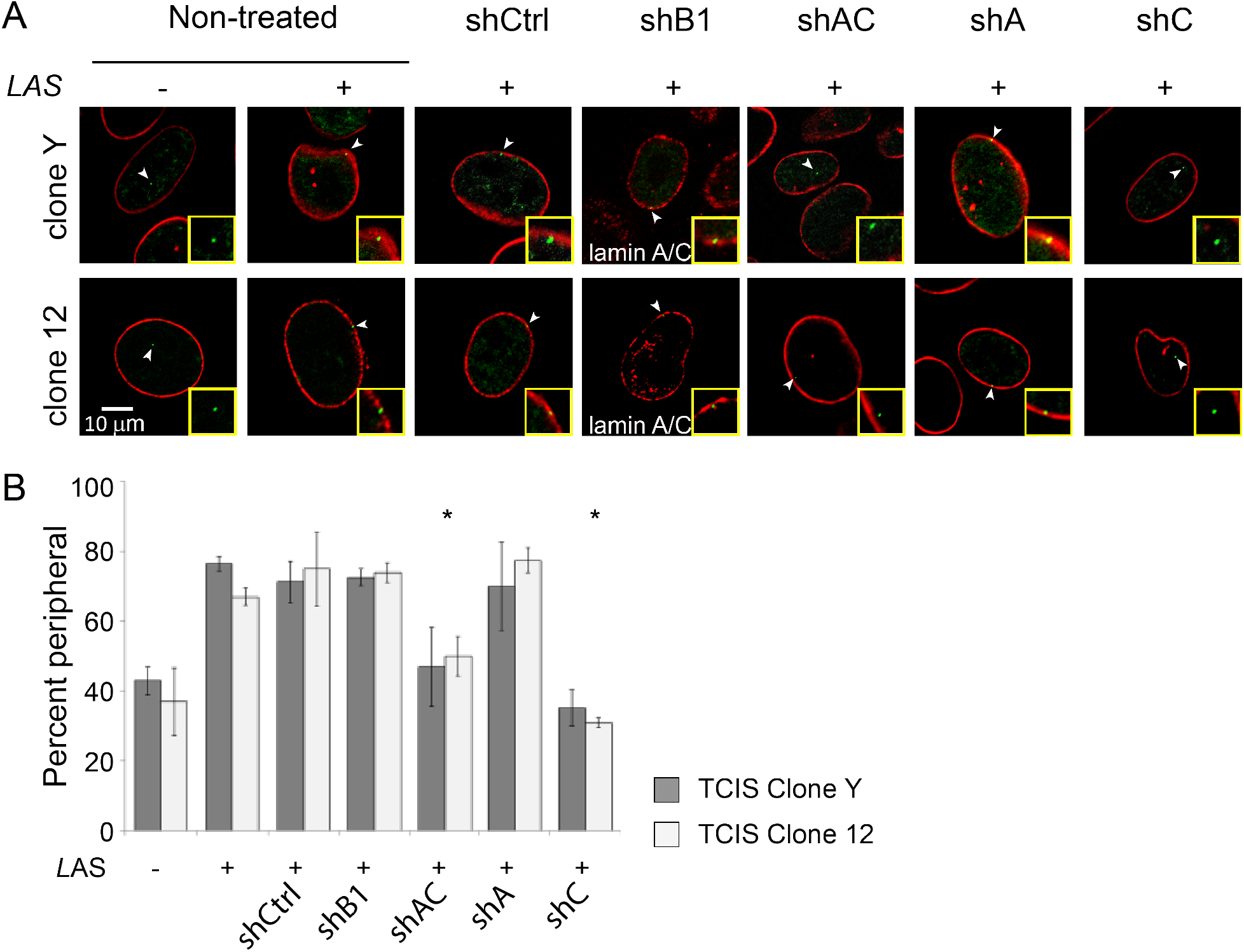
Lamin C is required for recruitment of chromatin to the lamina. (A) Representative images showing the disposition of *lacO* arrays (arrowheads, green) and lamin B1 (red, lamin A/C in shB1 treated cells) in the TCIS clones Y (top) and 12 (bottom) pre- and post- “switching” in of the Ikzf LAS and the effects of specific lamin depletion on the position of the *lacO* arrays. The inset shows 300× magnification. (B) Quantification of peripheral association was determined by overlap of EGFP-LacI foci and Lamin B1 (n ≥ 50). Error bars indicate SD. p ≤ 0.001 is indicated by asterisks.

We used this TCIS-LAS which had high association with the NE to monitor and quantify LAD organization in MEFs depleted of lamin isotypes. Removing lamin B1 alone (shB1) or lamin A alone (shA) had no significant effect (TCIS-LAS remained lamina-proximal), whereas NE-association was reduced significantly in cells depleted of both A and C (Fig. 2; shAC, p<0.001), or lamin C alone (Fig. 2B; shC, p<0.001). Without lamin C the percentage of NE-associated TCIS-LAS loci was reduced to the background seen with the LAS-less TCIS cassette (Fig. 2B). These data are consistent with our previous finding that acute shRNA-mediated removal of lamin A was insufficient to perturb LAS NE-association^6^. Given that shC treated cells showed loss of LAD organization, we next verified that this phenotype could be rescued by expression of mCherry-tagged lamin C, but not lamin A. We stably expressed either mCherry-tagged human lamin A or mCherry-tagged human lamin C in shC treated MEFs. Importantly, these constructs were not targeted by our murine-specific shRNA. TCIS-LAS localization at the NE was fully restored by mCherry human lamin C, and not by mCherry human lamin A (Fig. S9). These results independently support the hypothesis that lamin C, in contrast to lamin A and lamin B1, is required to maintain LAD association with the NE and nuclear lamina.

### Lamin C is nucleoplasmic during telophase and early G1-phase, and is significantly delayed in its association with the reforming NE

Our results thus far show that lamin C is important for normal interphase LAD configuration in MEFs (Fig. 1, 2). After mitosis, the nuclear lamina itself must be rebuilt and organized. The major consensus is that B type lamins associate with the nascent NE prior to A type lamins, and evidence of the differential dynamics of NE incorporation for lamin A versus lamin C are conflicting. In support of lamin C incorporating at the NE after lamin A, a study using injected recombinant proteins showed lamin A exhibited much faster lamina incorporation kinetics (20min) compared to lamin C (180 min)^47^. Even at 180min post-injection, nucleoplasmic lamin C foci were still evident. However, in that study, the incorporation of lamin C into the lamina was accelerated upon co-injection with lamin A, suggesting some cross regulation, in agreement with another study suggesting lamin C localization is dependent on lamin A^48^. A drawback of this study is that the normal regulation of both A/C ratios and post-translational modifications (PTM) are lacking and other studies have correct localization of lamin C to the NE in the absence of lamin A^33^. In support of lamin A and C arriving at the NE at the same time, a study using lamin A and lamin C over-expressed individually found that both A type lamins post-mitosis had similar kinetics of NE localization^49^. Of interest, interphase LAD and chromatin compartment organization is also ablated during mitosis and must also be re-built in the next G1, with overall chromosome positions and chromatin domains faithfully reinstated^8,9,16,24,25,30^. Yet, the pathway(s) and the mechanisms by which LADs reorganize and re-associate with the nuclear lamina after mitosis are still not understood.

In order to understand the role that lamin C might play in this process and given the conflicting data regarding timing of lamin C incorporation to the lamina, we first sought to evaluate post-mitotic lamin isotype NE incorporation dynamics in our MEFs. We imaged lamins A, C and B1 during exit from mitosis using a specific lamin B1 antibody in MEFs co-expressing mCherry-lamin A and EYFP-lamin C (Fig. 3). Localization of each lamin isotype was measured by fluorescence intensity histograms for a minimum of 20 nuclei along a line drawn through the medial plane of the nuclear volume as determined by Hoechst signal (Fig. 3). We found that all isotypes tested, lamins A, C and B1, concentrate at the NE during interphase (Fig. 3 and Fig. S10a, b), with lamin C and, to a lesser degree, lamin A also localizing diffusely in the nucleoplasm (Fig. 3 and Fig. S10a, b). Such nucleoplasmic localization was not unexpected given previous reports describing a ‘nuclear veil’ of lamin A/C^50–52^. However, during telophase, when mCherry-lamin A and lamin B1 were already highly enriched at or localized to the nascent NE, EYFP-lamin C was solely nucleoplasmic, with no detectable concentration at the NE (Fig. 3A, B and Fig. S11a, b). Similar results were obtained using a purportedly lamin C-specific antibody (Fig. S12). These results were highly reproducible from cell to cell (Fig. S11a, b), with EYFP-lamin C persisting predominantly in the nuclear interior well into early G1, and then gradually incorporating into the nuclear lamina.

**Figure 3:**
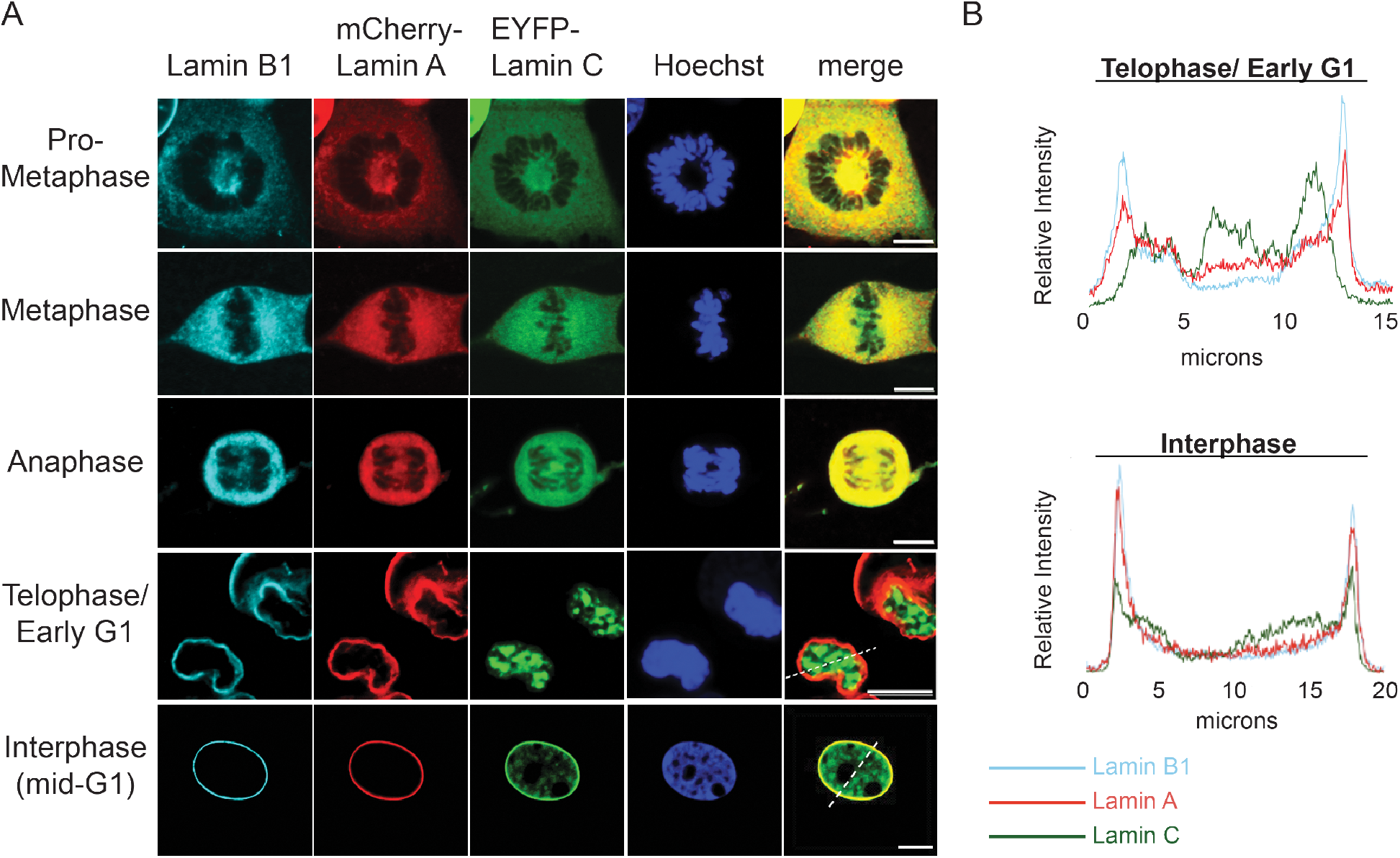
Lamin C persists in the nucleoplasm after mitosis. (A) Representative images of lamin B1 (cyan), lamin A (red), lamin C (green) and chromatin (blue) in different stages of the cell cycle. Merged images show lamins A and C localization. Dotted lines on the merged images for telophase/early G1 and G1 indicate the segment used for line scan displays shown in (B). (B) shows representative plots of intensities of lamin B1 (cyan), lamin A (red) and lamin C (green) along the dotted lines (from left to right) as shown in the merged images in (A) for the telophase/early G1 transition stage of the cell cycle and the mid-G1 (interphase) stage. Scale bar is 10*μ*m.

This delayed arrival of lamin C at the NE during early G1-phase was reminiscent of our previous finding that LADs are initially nucleoplasmic and exhibit delayed re-incorporation at the nuclear lamina after mitosis^9,16^. LADs themselves form nucleoplasmic intra-chromosomal LAD:LAD agglomerations during telophase and early-G1, and only later re-associate with the NE; as cells progress further into G1-phase, more LADs become NE-proximal and flattened against the lamina, with a subset of LADs remaining in the nuclear interior for up to three or four hours, well into G1-phase, quite reminiscent of the timing we noted for lamin C NE incorporation^9,16^. This led us to question whether lamin C might co-localize with LADs at the end of mitosis and into early G1. To test this we used a LAD-tracer system to fluorescently identify all endogenous LADs in living cells (MEFs) co-expressing EYFP-lamin C^9,16,53^. The LAD-tracer system relies on the expression of two constructs. First, a construct expressing Dam-lamin B1 enables methylation of DNA at adenine residues in proximity to the nuclear periphery (i.e. marks LADs with ^me^A), but has two additional domains that strictly control its expression: a destabilization domain (DD) that causes degradation in the absence of the shield ligand, and a Cdc10 dependent transcript 1(CDT) regulatory domain restricts its expression to G1^9,16,53,54^. The second construct, the LAD-tracer, is a modified mCherry-tagged version of the previously described m6A-tracer^16,53^ that binds ^me^A, the modification generated by Dam-Lamin B1, thus marking LADs with mCherry. The LAD-tracer is expressed throughout the cell cycle and identifies ^me^A-modified DNA (LADs) in all phases in cells where both constructs are expressed at appropriate levels. As expected, EYFP-lamin C co-localized with the LAD-tracer in interphase cells (Fig. 4A, S13, movie 1, and movie 2). However, during telophase and early-G1 we were surprised to find that lamin C and LADs occupied distinct nuclear volumes with minimal or no co-localization (Fig. 4A, B and S14). Despite their apparent separation, time-lapse movies showed that EYFP-lamin C and LAD aggregates had coincident arrival to and integration with the nuclear lamina (Fig. 4C, S15, movie 3, and movie 4). These results suggest lamin C functions distal to LADs to influence their organization.

**Figure 4:**
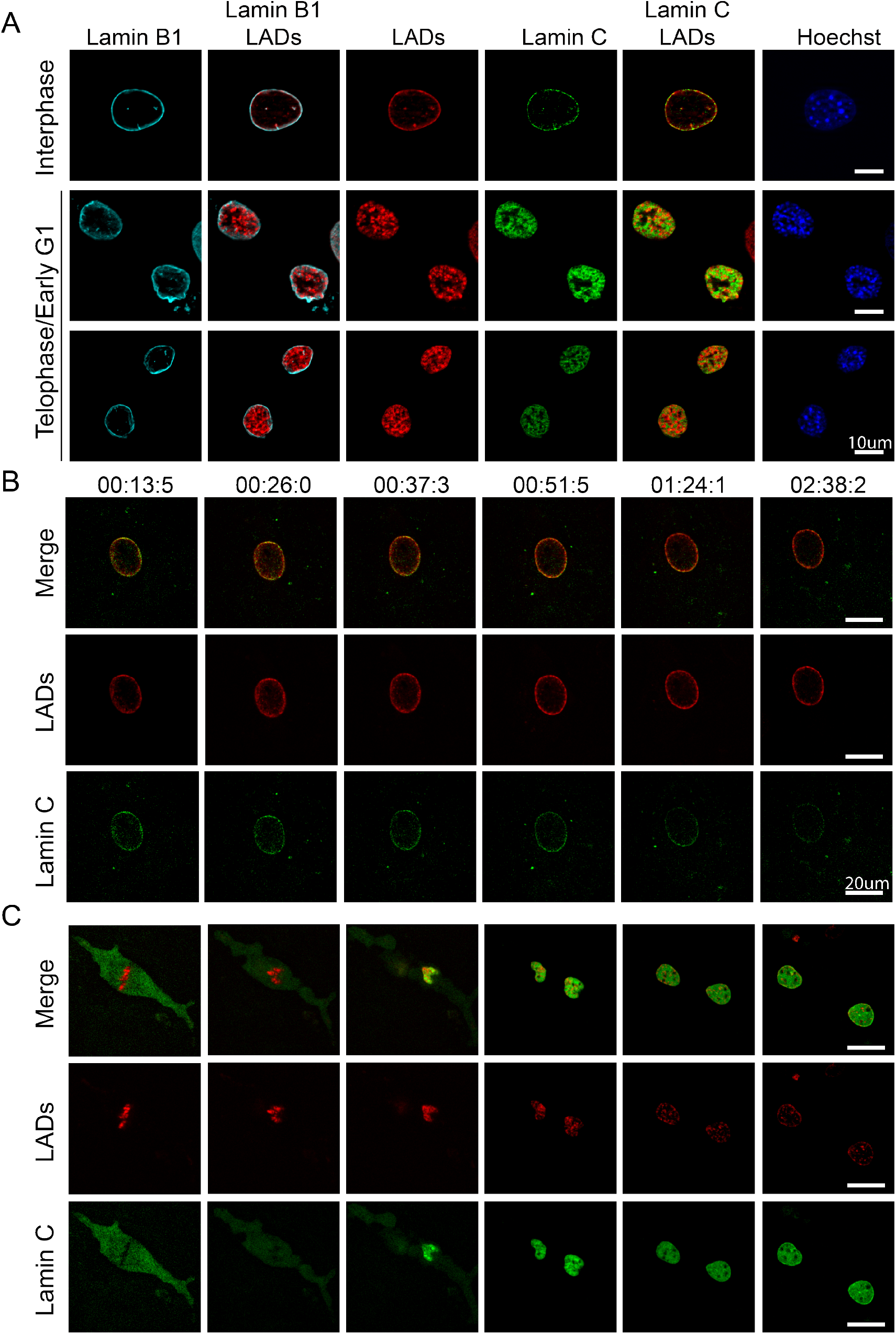
Lamin C and LADs are recruited to the NE in G1, but are not co-localized in the nucleoplasm. (A)Representative images of interphase and early G1 nuclei with anti-lamin B1 (cyan), LAD-tracer (red), and lamin C (green), DNA (blue). (B) Still images from time-lapse movie 1 of LADs (red) EYFP-lamin C (green) during interphase. Scale bar is 20*μ*m (C). Still images from time lapse movie 3 of LADs (red) EYFP-Lamin C (green) during mitosis. Scale bar is 20*μ*m. Images were chosen to exemplify certain stages (metaphase, anaphase, telophase, early G1, partially resolved, fully resolved).

### Lamin C is required during early G1 for LAD integrity and LAD recruitment to the NE

To test if lamin C might play a role in restoring LAD organization after mitosis, we used shRNAs to specifically deplete lamin A or lamin C for four days to allow lamin turnover in MEF cells harboring the LAD-tracer system. These shRNA-treated cells were then subjected to a single thymidine block (24 hours), followed by release into enriched media for several hours and subsequently treated with the Cdk1 inhibitor RO3306 to arrest them at the G2/M transition^9,55^. After overnight block, cells were released into complete medium (without shield ligand). Cells rapidly entered mitosis and were examined at subsequent time points, up to four hours later. Because LAD targeting to the NE after mitosis is normally gradual, over several hours, we chose to assay cells four hours after release from the G2/M block, when a majority of control cells have exited mitosis and show ‘resolved’ interphase LAD organization (Fig. 5A). Nuclei were independently scored by two observers as either ‘resolved’ (all LAD-tracer signal adjacent to lamin B1 NE signal) or ‘unresolved’ (if any LAD-tracer signal was not adjacent to lamin B1); nuclei were counted over groups of 5 frames and then averaged across groups of 5 (n=4 groups, >150 nuclei). Nuclei lacking LAD-tracer signal were not scored. For the shCtrl-treated cells, 44% of nuclei had unresolved LADs. We note that this background of ‘unresolved’ LADs is mainly due to timing and shortcomings inherent in systems requiring co-expression of two independent components. We estimated that about 30% of LAD-tracer expressing nuclei simply under-express Dam-lamin B1 relative to LAD-tracer, leading to accumulation of diffuse mCherry in the nucleoplasm even in Mid-G1 and, while qualitatively different, these were included in the ‘unresolved’ numbers. Cells depleted of lamin A showed a similar background, with 38% unresolved nuclei (shA; Fig. 5), again suggesting lamin A has no active role in post-mitotic LAD assembly at the NE. By contrast, in the lamin C knockdown population, 62% of nuclei had unresolved LADs, a significant increase of nearly 50% over controls (Fig. 5; p<0.001). Interestingly, many disrupted lamin C-depleted nuclei revealed an additional phenotype: LAD aggregates appeared to have decondensed slightly, sometimes forming string-like networks in the nucleoplasm (Fig. 5A), quite distinct from the compact NE-associated LADs in controls (Fig. S16). This conformation of the heterochromatic LADs was even visible via Hoechst stain (Fig. 5A). Such string-like networks were absent from control and shA treated cells and differed from LAD organization seen in untreated or lamin A-depleted cells immediately after mitosis, where LADs not at the NE remained in clear condensed and separated domains. It is important to note that at 4 hours post release, the percentages of cells in M, early G1 and mid-G1/S/G2 were quite similar for all treatments. However, two hours after release, while the percentages of cells in mitosis and early G1 were the same in control (shLacZ) and shC-treated cells, we note that shA and shAC treated cells displayed an apparent lag in early G1 at two hours (Fig. S1C). We indicate that this is an apparent lag since we quantify ‘early G1’ on morphometric measures (nuclear size, shape, and/or obvious match to a sister nucleus) and both nuclear shape and size could be influenced by loss of lamin A, since lamin A has been implicated in the mechanoregulation of nuclear morphology. The immediate post-mitosis stage, where the nuclei are still rounded up and the cytoskeleton has not yet exerted its influence on nuclear shape may be a particularly relevant point in the cell cycle for lamin A. This finding makes it even more striking that shA treated cells do not display perturbed lamin organization. Together, these findings collectively demonstrate that lamin C is required for LAD integrity and dynamic LAD recruitment and association with the NE and nuclear lamina after cell division.

**Figure 5:**
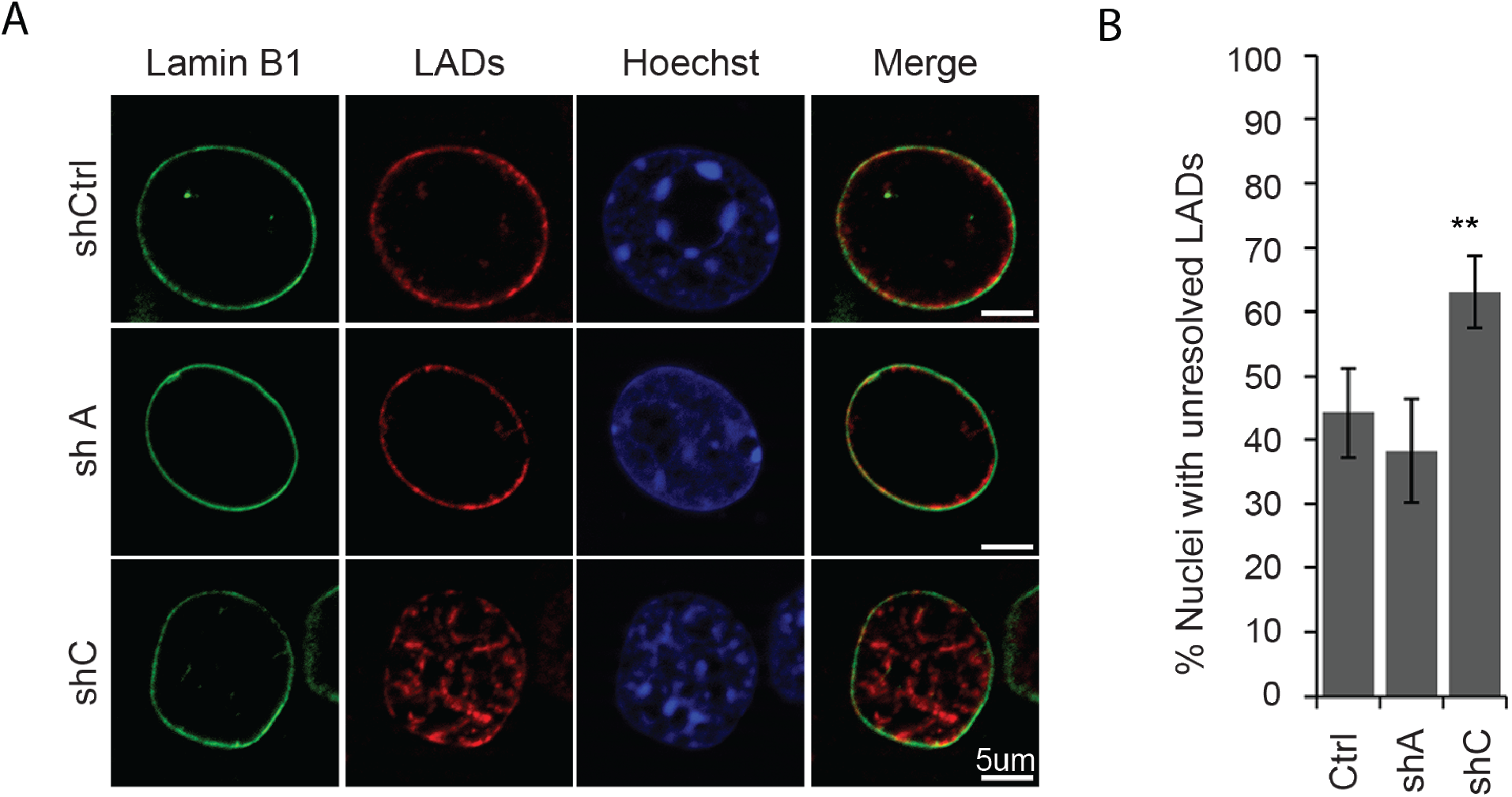
Depletion of lamin C leads to aberrant LAD accretion and localization to the NE. (A) Cells shown 4 hours after release from G2/M border. Anti-lamin B1 (green) LAD-tracer (red) Hoechst (blue) (B) Quantification (blind analysis) of nuclei with unresolved LADs (nuclei with LADs not at the periphery) 4 hours after release from the G2/M border (n ≥ 120). Error bars indicate SD. p ≤ 0.001 is indicated by asterisks

## Discussion

Our findings provide novel insights into 3D genome organization by showing that LAD integrity and post-mitotic association with the NE depend on lamin C, and not on lamin A. To examine the effects of removing specific A type lamin spliceoforms on LADs and overall genome organization, we employed three imaging approaches: (1) Chromosome conformation paints (CCP), (2) Tagged Chromosomal Insertion System (TCIS), and (3) the LAD-tracer system. These three methods enable examination of genome architecture at different levels of spatial and time resolution in both normal and lamin depleted cells. In addition, these methods provide single cell metrics of organization missed by bulk DamID analyses. Our CCP differentially label LADs and non-LADs across an entire chromosome, enabling us to visualize the organization of LADs and non-LADs, relative to each other and the nuclear lamina, in the context of the entire chromosome in situ. The TCIS system, on the other hand, allows us to observe changes to the peripheral localization of a single lamina associated genomic locus (LAS) and enables more robust quantification of perturbations to lamina association through a binary measure of lamina localization. Finally, the LAD-tracer system, which tags all LADs within the nucleus, allows us to measure the dynamics of LAD organization across the genome and relative to the nuclear lamina.

While we and others have previously implicated A type lamins in regulating LAD organization, we speculated that lamin A and lamin C might have different roles in LAD organization^6,44^. Both lamin A and lamin C are encoded by *LMNA* and cross-talk in their expression levels in a given cell type has been noted^47^. Several lines of evidence, from super-resolution microscopy to ectopic expression studies, indicate that lamin C and lamin A form unique networks that are nonetheless inter-dependent on some level^33–35^. Lamin C depleted mice (from birth) show little overt phenotype, with the exception of perturbed nuclear shapes, while mice expressing only lamin C (depleted of A) have longer lifespans^39–41^. Taken together, these data suggest that lamin A and C do indeed have differential roles in the nucleus. One such difference is in how lamin A and C form networks at the lamina, both in timing of association with the NE and in protein:protein interactions. For instance, lamin C preferentially interacts with nuclear pore complexes and altered A/C ratios change the mechanical properties of the nucleus^33,56^. In this study, we identified a critical role for lamin C in genome organization. In particular, lamin C is critical for both organizing LADs to the nuclear lamina and for LAD sub-chromosomal domain integrity, since acute depletion of this isotype caused derangement and inter-mixing of LAD/non-LAD (A/B compartment) chromatin (Fig. 1).

Interphase genome organization, including LAD and lamin organization, is ablated during mitosis and re-established in the subsequent G1 phase^8,9,16,24,25,28–30^. Previous studies suggest that A-type lamins organize to the reforming nuclear envelope with different kinetics, although there is some discrepancy on how lamin A and C might differ in their timing of association^47–49,56^. We find, in agreement with previous studies, that lamin B1 incorporates into the reforming NE at anaphase, preceding both lamin A and C incorporation. Our results, using both fluorescent proteins as well as immunofluorescence, indicate that lamin A associates with the NE prior to lamin C in MEFs, with the majority of lamin C remaining nucleoplasmic well into early G1 (Fig. 3, S11a, b). This is intriguing given that lamin C appears to be critical for LAD organization (Fig. 1 and 2) and we had previously shown that LADs are also nucleoplasmic during this stage of the cell cycle^9^.

To further define the spatial and temporal relationship between LAD organization and lamin C organization during the critical transition from mitosis into G1, we used the LAD-tracer system to demarcate LADs in EYFP-lamin C. Strikingly, lamin C was excluded from the heterochromatic LADs at mitotic exit well into early G1 (3 hours), suggesting that lamin C predominantly interacts with euchromatic regions of the genome or is excluded from heterochromatic regions in early G1 (Fig. 4, S13, S14, movie 3 and movie 4). Overall our data suggests that the A-type lamin isotype bound to euchromatic regions is predominantly lamin C, which we find to be present at higher levels relative to lamin A in the nucleoplasm where euchromatin is enriched in interphase. We suggest that these interactions are established at the end of mitosis and persist into interphase (See Fig. S11). Our conclusion that lamin C co-localizes with euchromatic regions at mitotic exit is particularly intriguing given a recent study that found that interphase nucleoplasmic A-type lamins, and in particular lamin C, preferentially interact with enhancers and promoters of active genes^52^. The lamins bound to these euchromatic regions remain phosphorylated on Serine 22 (pS22, a known mitotic PTM) even after mitotic exit. Intriguingly, the sites at which pS22-lamin C binds is altered in Hutchinson Gilford Progeria Syndrome (HGPS) and is correlated with up-regulation of genes. Previous studies have also identified nucleoplasmic lamins bound to euchromatic regions that are reliant on the nucleoplasmic LAP2 (lamina associated peptide 2) isoform LAP2*α* ^57,58^. It is unknown how the interactions between lamin C and LAP2*α* are mediated, but such associations could be the result of PTM regulation on A type lamins. In addition, as demonstrated by altered pS22- lamin C chromatin binding and gene activation in HGPS, which results from a lamin A specific mutation, the interdependence of the lamin networks in regulating gene activity (and other processes) is likely an important aspect of these diseases. Finally, a previous proteomics study found that lamin C preferentially interacts with components of the NPC, a complex associated with euchromatin and depleted in heterochromatin^33^. Taken together, these studies and our new results describe a role for lamin C independent of lamin A and acting through interactions with the euchromatic compartment of the genome.

Thus, as important as lamin C is for organization of LADs, the post-mitotic organization of these heterochromatic domains to the lamina via guided transit towards the NE through direct interactions with lamin C, is unlikely. What role might lamin C be playing during the transition from mitosis into G1? If lamin C is not directly interacting with LADs, how can it be such an important regulator of LAD organization? We find that, in the absence of lamin C (but not lamin A), LAD aggregates are delayed or prevented in their association with the NE. Importantly, the LADs appear to form string-like networks of interconnected aggregates (Fig. 5), strongly suggesting that there is a problem in spatially segregating the forming LAD/non-LAD (A/B) chromatin compartments. This is supported by our CCP studies (Fig. 1), in which we observe gross disruption of spatial organization of LAD:LAD interactions, loss of clear A/B intra-chromosomal domain organization, and loss of LAD association with the NE. Our findings therefore suggest that lamin C is dynamically spatially regulated during exit from mitosis to promote novel associations needed for LAD control, and/or to block aberrant interactions and premature reassembly at the NE.

### Speculations and a model

We postulate that the temporal separation of incorporation of lamin isotypes at the nuclear periphery post mitosis allows for the formation of separate, but interdependent, lamina meshworks, as has been reported^33^. Our data suggests that lamin B1 NE meshworks are formed first, followed by lamin A and, lastly, lamin C, much of which remains nucleoplasmic throughout interphase. We speculate that the Serine 22 phosphorylation blocks or alters polymerization of A-type lamins, and enables a control in timing and level of incorporation into the NE^35,48,52^. The late recruitment of lamin C to the NE post-mitosis is strikingly coincident with LAD accumulation to this region, suggesting a coordinated regulation which is supported by LAD disruption in the absence of lamin C. We further speculate that, prior to its accumulation at the nascent NE, lamin C surrounds but is excluded from LADs and might ‘instruct’ genome reorganization by promoting robust LAD-LAD self-association within each chromosome and preventing LAD aggregation between chromosomes; a potential danger since euchromatin and heterochromatin are each capable of self-aggregating via phase partitioning^22^. Without a mechanism to prevent wide-spread self-aggregation (or accretion), heterochromatin might globally cluster, leading to chromosome entanglement and difficulty accessing appropriate genes. In support of this, regulated wholesale agglomeration of heterochromatin in the center of the nucleus is reported in cell types that lack A-type lamins and LBR, such as rod photoreceptor cells^59^. We therefore propose a gross overall schema (Fig. 6) outlining a suggested role for lamin C in the reorganization of the genome post-mitosis. We propose that lamin C might directly associate with euchromatin or interact with chromosome scaffolding proteins (such as CTCF) which are enriched on promoters, enhancers, and borders of compartments and LADs^9,14^. From a recent study, we propose that lamin C with phosphorylated Ser22, a mitotic modification, will be retained in the nucleoplasm and interact with euchromatin, potentially through its interactions with LAP2*α* or NPC components^52^. In this model, a heterochromatin (LAD) core would be surrounded by lamin C monomers or short polymers which would serve as a ‘buffer’ between chromosomes. These interactions would also reinforce the separation between A- and B-compartment chromatin in each chromosome and prevent aberrant ‘sticky’ heterochromatin interactions between chromosomes. Lastly, this dual purposed segregation mechanism persists as both LADs and lamin C accumulate at the nuclear periphery, with the latter potentially dependent on or enriched in interactions with NPCs and their associated underlying euchromatin^60,61^.

**Figure 6:**
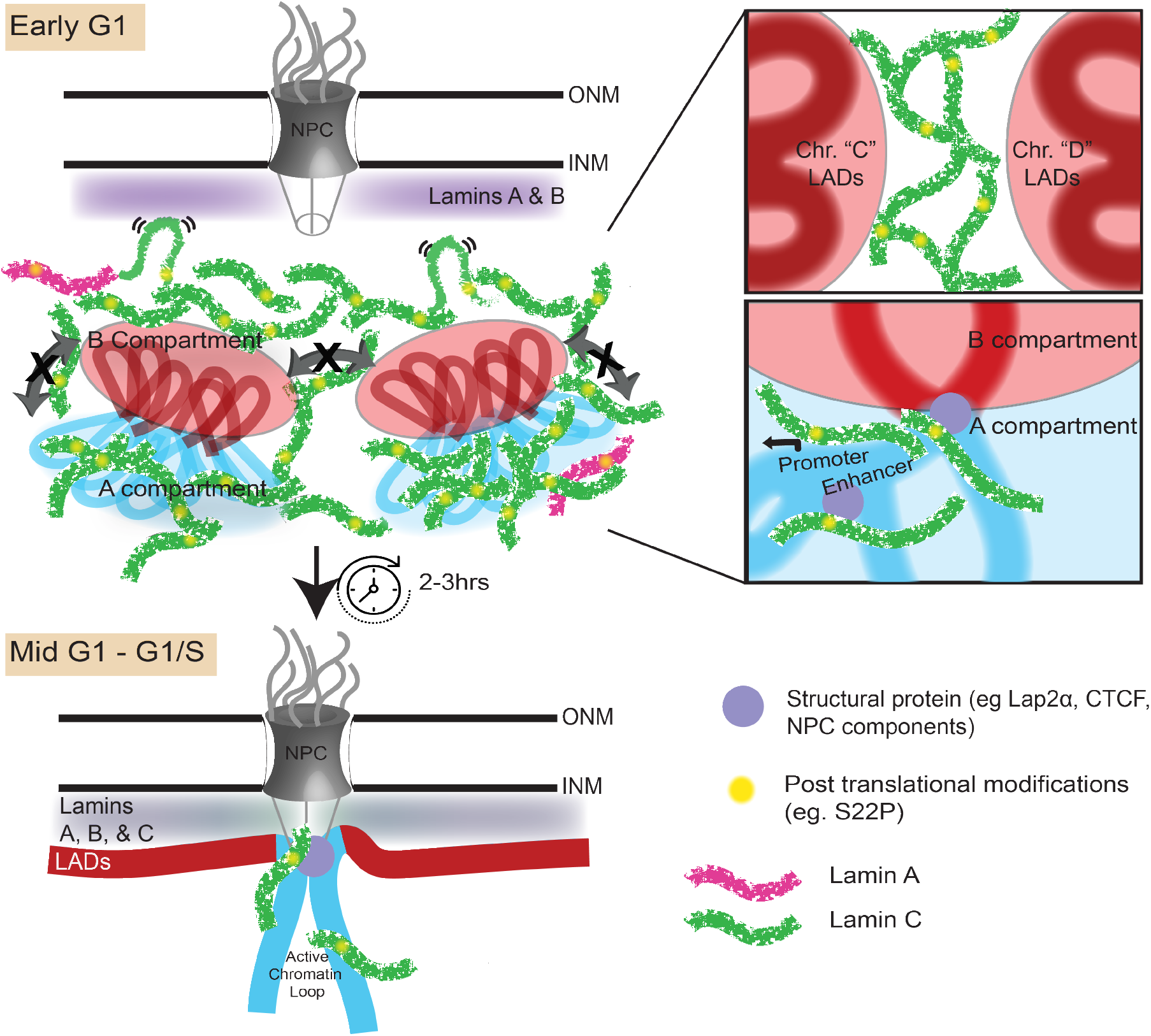
Speculative model of the role of lamin C in genome organization. Post translational modifications (e.g. Phosphoserine 22) of lamin C allows its nucleoplasmic localization during mitotic exit and into early G1. During this phase lamin C is spatially excluded from LADs potentially by binding to structural proteins such as CTCF, LAP2*α* etc on euchromatin and/or phase separation phenomena thereby physically hindering aberrant inter-chromosomal LAD interactions (Top inset) and reinforcing intra-chromosomal A-B compartmentalization (Bottom inset).

In summary, we discovered that lamin C is uniquely required to efficiently target LADs to the NE after cell division and to maintain the integrity of A/B compartments and overall gross 3D genome organization. We propose that lamin C promotes intra-chromosomal LAD aggregation and prevents aberrant trans-chromosomal heterochromatin interactions. Our results bring up several additional questions, including the role these proteins have in organizing and regulating the genome during development and in laminopathies, particularly those in which lamin C is not directly disrupted, but where its deregulation or mis-localization might contribute to pathology. To fully understand the molecular pathway by which cells re-establish tissue-specific 3D genome architecture after mitosis, it will be important to establish both the role of post-mitotic PTMs of lamin C and the lamin C protein interactome during exit from mitosis.

## Materials and Methods

### DamID-seq datasets

The data discussed in this publication have been deposited in NCBI’s Gene Expression Omnibus (Edgar et al., 2002) and are accessible through GEO Series accession number GSE97095 (https://www.ncbi.nlm.nih.gov/geo/query/acc.cgi?acc=GSE97095).

### Contact for reagent and resource sharing

Further information and requests for resources and reagents should be directed to and will be fulfilled by the Lead Contact, Karen Reddy.

### Generation and maintenance of primary murine embryonic fibroblast (MEFs)

For primary MEFs, wild-type eight-week-old C57BL/6J mice were bred and embryos were harvested at E13.5. Individual embryos were homogenized using a razor blade, and cells were dissociated in 3 ml 0.05% trypsin for 20 min at 37°C, then 2 ml of 0.25% trypsin was added and incubated again at 37°C for 5 min. Cells were pipetted vigorously to establish single cells, passed through a 70 *μ*m cell strainer, pelleted and then plated in 10 cm dishes and labeled as P0. MEFs were cultured in DMEM High Glucose with 10% FBS, penicillin/streptomycin, L-glutamine and non-essential amino acids. Cells were cultured for no longer than 5 passages before harvesting for experiments. For initial DamID and m6A tracer experiments, longer term-culture C57BL/6 MEFs were purchased from ATCC (American Tissue Culture Collection, CRL-2752) and cultured according to their established protocols, in medium containing DMEM High, 10% FBS, Penicillin/Streptomycin and L-glutamine.

### Lamin A/C knockdown

shRNA-mediated knockdown was carried out as described previously. Specifically, virus for knockdowns were generated in HEK 293T/17 cells (ATCC CRL-11268) by co-transfecting VSV-G, delta 8.9, and a plko.1 vector driving the expression of control shRNAs - shluciferase (5′-CGCTGAGTACTTCGAAATGTC-3′) or shLacz (5′-CGCTAAATACTGGCAGGCGTT-3′), shLmnA/C(Sigma clone NM_001002011.2-901s21c, 5′-GCGGCTTGTGGAGATCGATAA-3′), shLmnA (produced in our lab), shLMNC (produced in our lab, 5′-TCTCCCACCTCCATGCCAAAG-3’) or shLMNB1 (Sigma clone NM_010721.1-956s1c1, 5′-GCGTCAGATTGAGTATGAGTA-3′) with Fugene 6 transfection reagent (Promega E2691). 10 mM sodium butyrate was then added to the transfected cells 3 hours post transfection for an overnight incubation at 37°C, 5% CO_2_. The transfection media containing sodium butyrate was removed the following day and the cells were washed with 1X PBS. Opti-MEM was then added back to the cells which were then incubated at 37°C, 5% CO_2_. Viral supernatant was collected every 12 hours up to 3 collections and the supernatant of all 3 collections were pooled. Primary MEFs were cultured as described and incubated overnight with different shRNA viruses per condition supplemented with 4 *μ*g/ml polybrene and 10% FBS for 12-14 hours. Fresh MEF media was then added to the cells after the virus was removed and selected with 20 *μ*g/ml blasticidin or 2 *μ*g/ml puromycin. For DamID profiling, cells were infected with DamID virus 4 days post shRNA transduction and cultured for an additional 48 hours.

### DamID Infection

DamID was performed as described previously^5,6,14,62,63^. Cells were transduced with lentiviruses harboring the Dam constructs. Lentiviral vectors pLGW-Dam and pLGW Dam-LmnB1 were co-transfected with VSV-G and delta 8.9 into HEK 293T/17 packaging cells using the Fugene 6 transfection reagent in DMEM High glucose complete media (DMEM High glucose supplemented with 10% FBS, Penicillin/Streptomycin, L-glutamine). 10 mM sodium butyrate was added to the transfected cells 3 hours post-transfection and left overnight. The following day this media was removed and the cells were washed briefly with 1X PBS before Opti-MEM media was added. Supernatants containing viral particles were collected every 12 hours between 36-72 hours after transfection, and these collections were pooled, filtered through 0.45 *μ*M SFCA or PES, and then concentrated by ultracentrifugation. For infection with lentivirus, MEFs were incubated overnight with either Dam-only or Dam-LmnB1 viral supernatant and 4 *μ*g polybrene. Cells were allowed to expand for 2-4 days then pelleted for harvest.

### DamID protocol

MEFs were collected by trypsinization and DNA was isolated using QIAamp DNA Mini kit (Qiagen, 51304), followed by ethanol precipitation and resuspension to 1 *μ*g/*μ*l in 10 mM Tris, pH 8.0. Digestion was performed overnight using 0.5-2.5 *μ*g of this genomic DNA and restriction enzyme DpnI (NEB, R0176) and then heat-killed for 20 minutes at 80°C. Samples were cooled, then double stranded adapters of annealed oligonucleotides (IDT, HPLC purified) AdRt (5′-CTAATACGACTCACTATAGGGCAGCGTGGTCGCGGCCGAGGA-3′) and AdRb (5′-TCCTCGGCCG-3′) were ligated to the DpnI digested fragments in an overnight reaction at 16° C using T4 DNA ligase (Roche, 799009). After incubation the ligase was heat-inactivated at 65° C for 10 minutes, samples were cooled and then digested with DpnII for one hour at 37° C (NEB, R0543). These ligated pools were then amplified using AdR_PCR oligonucleotides as primer (5′-GGTCGCGGCCGAGGATC-3′) (IDT) and Advantage cDNA polymerase mix (Clontech, 639105). Amplicons were electrophoresed in 1% agarose gel to check for amplification and the size distribution of the library and then column purified (Qiagen, 28104). Once purified, material was checked for LAD enrichment via qPCR (Applied Biosystems, 4368577 and StepOne Plus machine) using controls specific to an internal Immunoglobulin heavy chain (*Igh)* LAD region (J558 1, 5′-AGTGCAGGGCTCACAGAAAA-3′, and J558 12, 5′-CAGCTCCATCCCATGGTTAGA-3′) for validation prior to sequencing.

### DamID-seq Library Preparation

In order to ensure sequencing of all DamID fragments, post-DamID amplified material was randomized by performing an end repair reaction, followed by ligation and sonication. Specifically, 0.5-5 *μ*g of column purified DamID material (from above) was end-repaired using the NEBNext End Repair Module (NEB E6050S) following manufacturer’s recommendations. After purification using the QIAquick PCR Purification Kit (Qiagen, 28104), 1*μ*g of this material was then ligated in a volume of 20 *μ*l with 1 *μ*l of T4 DNA ligase (Roche, 10799009001) at 16°C to generate a randomized library of large fragments. These large fragments were sonicated (in a volume of 200μl, 10mM Tris, pH 8.0) to generate fragments suitable for sequencing using a Bioruptor^®^ UCD-200 at high power, 30 seconds ON, 30 seconds OFF for 1 hour in a 1.5 mL DNA LoBind microfuge tube (Eppendorf, 022431005). The DNA was then transferred to 1.5 ml TPX tubes (Diagenode, C30010010-1000) and sonicated for 4 rounds of 10 minutes (high power, 30 seconds ON and 30 seconds OFF). The DNA was transferred to new TPX tubes after each round to prevent etching of the TPX plastic. The sonication procedure yielded DNA sizes ranging from 100-200 bp. After sonication, the DNA was precipitated by adding 20 *μ*l of 3M sodium acetate pH 5.5, 500 *μ*l ethanol and supplemented with 3 *μ*l of glycogen (molecular biology grade, 20 mg/ml) and kept at −80°C for at least 2 hours. The DNA mix was centrifuged at full speed for 10 min to pellet the sheared DNA with the carrier glycogen. The pellet was washed with 70% ethanol and then centrifuged again at full speed. The DNA pellet was then left to air dry. 20 *μ*l of 10 mM Tris-HCl was used to resuspend the DNA pellet. 1 *μ*l was quantified using the Quant-iT PicoGreen dsDNA kit (Invitrogen, P7589). Sequencing library preparation was performed using the NEBNext Ultra DNA library prep kit for Illumina (NEB, E7370S), following manufacturer instructions. Library quality and size was determined using a Bioanalyzer 2100 with DNA High Sensitivity reagents (Agilent, 5067-4626). Libraries were then quantified using the Kapa quantification Complete kit for Illumina (Kapa Biosystems, KK4824) on an Applied Biosystems 7500 Real Time qPCR system. Samples were normalized and pooled for multiplex sequencing.

### DamID-seq data processing

DamID-seq reads were processed using LADetector (https://github.com/thereddylab/pyLAD), an updated and packaged version of the circular binary segmentation strategy previously described for identifying LADs from either array or sequencing data (https://github.com/thereddylab/LADetector)^5,6^. LADs separated by less than 25 kb were considered to be part of a single LAD. All other parameters were left at default values. LADs were post-filtered to be greater than 100 kb, complementary genomic regions to LADs were defined as non-LADs. Bed files were generated for visualization using the pyLAD LADetector.

### LAD and non-LAD chromosome-wide probe design and labeling

LADs from murine embryonic fibroblasts were defined through the LADetector algorithm, and complementary regions to Chromosomes 11 and 12 were defined as non-LADs^9,16,53,54^. Data provided Geo GSE56990. Centromeres were excluded, and LAD and non-LADs were repeat masked. Probes were selected in silico based on TM and GC content, and those with high homology to off target loci were specifically removed. 150 base pair oligos were chemically synthesized using proprietary Agilent technology and probes were labeled with either Cy3 or Cy5 dyes using the Genomic DNA ULS Labeling Kit (Agilent, 5190-0419). 40 ng of LAD and non-LAD probes were combined with hybridization solution (10% dextran sulfate, 50% formamide, 2X SSC) then denatured at 98°C for 5 minutes and pre-annealed at 37°C.

### 3D-ImmunoFISH and immunofluorescence

3D-immunoFISH was performed as described previously^6,63^. Briefly, primary mouse embryonic fibroblast cells were plated on poly-L-lysine coated slides overnight. Cells on slides were fixed in 4% paraformaldehyde (PFA)/1X PBS for 16 minutes, then subjected to 3-5 minute washes in 1X PBS. After fixation and washing, cells were permeabilized in 0.5% TritonX-100/0.5% saponin for 15-20 minutes. The cells were washed 3 times 5 minutes each wash in 1X PBS, then acid treated in 0.1N hydrochloric acid for 12 minutes at room temperature. After acid treatment, slides were placed directly in 20% glycerol/1X PBS and then incubated at least one hour at room temperature or overnight at 4° C. After soaking in glycerol, cells were subjected to 4 freeze/thaw cycles by immersing glycerol coated slides in a liquid nitrogen bath. Cells were treated with RNAse (100 *μ*g/ml) for 15 min in 2X SSC at room temperature in a humidified chamber. DNA in cells was denatured by incubating the slides in 70% formamide/2X SSC at 74° C for 3 min, then 50% formamide/2X SSC at 74° C for 1 min. After this denaturation, cells were covered with a coverslip containing chromosome conformation paints in hybridization solution and sealed. After overnight incubation at 37° C, slides were washed three times in 50% formamide/2X SSC at 47° C, three times with 63° C 0.2X SSC, one time with 2X SSC, and then two times with 1X PBS before blocking with 4% BSA in PBS for 30-60 min in a humidified chamber. Slides were then incubated with anti-LmnB1 primary antibody (1:200 dilution; Santa Cruz, SC-6217) in blocking medium overnight at 4° C. Slides were washed three times with 1X PBS/0.05% Triton X-100 and then incubated with secondary antibody in blocking medium Alexa Fluor 488 (1:200 dilution; A32814) for 1 hour at room temperature. Post incubation, slides were washed three times with 1X PBS/0.05% Triton X-100, and then DNA counterstained with 1 *μ*g/ml Hoechst. Slides were then washed, mounted with SlowFade Gold (Life Technologies, S36936).

### Live Cell imaging

Immortalized C57Bl/6 MEFs (ATCC CRL-2752) cells were infected to stably express ddDam-LaminB1-CDT, EYFP-Lamin C and m6A tracer. ddDam-LaminB1-CDT is a destabilized version of the previously described DamID construct that has incorporated the CDT domain from the Fucci system to ensure its expression is restricted to interphase^9,16,53,54^. The m6A-tracer is comprised of a catalytically inactive version of DpnI, that retains its ability to bind DNA, in frame with an mCherry red fluorescent protein^16,53^. For cell cycle experiments, these cells were grown in the presence of shield ligand (AOBIOUS, AOB6677), which stabilizes the ddDam-LaminB1-CDT, along with 1mM thymidine (Sigma) block for 24 hours to enable synchronization of cells at G1/S. This arrest was followed by release into complete DMEM medium (DMEM high glucose, +10% FBS, 100 U/mL Penicillin and 100 *μ*g/ml Streptomycin) containing 25*μ* M 2′-Deoxycytidine for 4 hours. Cells were then blocked at G2/M by incubation by replacing media with complete media containing 10uM R0-3306 (AOBIOUS, AOB2010) for 16-20 hours^55^. Cells were released from this block by washing 3 times with warm Fluorobrite DMEM +10% FBS with 100 U/mL Penicillin and 100 *μ*g/ml Streptomycin. 1-4 hours after release cells were imaged live every 1-5 using a 3i spinning disc confocal microscope. Interphase cells were not synchronized and were imaged every 1-5 minutes.

### TCIS

The two TCIS clones, clone Y and clone 12, harboring a lamina associated sequence (LAS) corresponding to a fragment of the Ikzf1 gene was generated as previously described^5,6^. C57BL/6 fibroblasts were transfected, using Fugene 6 (Promega), with a linearized TCIS construct described previously^6^. Cells were selected for hygromycin resistance (500 *μ*g/ml), and clones were isolated and expanded. Single integration clones were screened for by qPCR and transfection with EGFP-LacI retroviral vector to visualize the insert site. Clones 12 and Y had single integrations of the TCIS system at a chromosomal position away from the nuclear lamina, as determined by microscopy and either the presence or lack of an overlap in lamin B1 and EGFP-LacI accumulation at the *lacO*insert site. Site-specific recombination was obtained by co-transfection of TCIS clones with a DNA fragment corresponding to the LAS (Ikzf1) cloned into a switch vector and Cre recombinase. Switched cells were then seeded at low density with 10,000 cells per well of a 6-well tissue culture dish and treated with 1 *μ*M ganciclovir for 24 h. TCIS cells require a short treatment with ganciclovir and to be treated at low confluence. Negative ganciclovir selection occurs when the non-switched thymidine kinase gene cassette expresses thymidine kinase, which in turn phosphorylates ganciclovir. Phosphorylated ganciclovir is toxic to the cells. Once released into the media, it can affect neighboring cells if not maintained at low confluence and if media is removed after 24 h. Cells that have successfully switched cannot phosphorylate ganciclovir and are therefore resistant. Cells resistant to ganciclovir (1 μM) were then expanded for nuclear positioning analysis. Transfections for specific recombination in TCIS clones were performed with the electroporation system (Amaxa Nucleofector 4; Lonza), to ensure essentially 100% transfection efficiency. Ingenio Electroporation Products (MIR 50111; Mirus Bio LLC) were used in combination with the Amaxa nucleofector. All cell lines were maintained in DMEM high with 10% FBS (U.S. Defined Fetal Bovine Serum; Hyclone) in the presence of 500 *μ*g/ml hygromycin (50 mg/ml; Hygromycin B; Corning/CellGro) and 1 mM IPTG when EGFP-LacI was present. To enable binding of EGFP-LacI, IPTG was removed from the cultures, and cells were analyzed after 24–36 h in fresh media.

### Fluorescent tagged lamin expression

Lentiviral vectors containing fluorescently tagged lamin A or C were co-transfected with VSV-G and delta 8.9 into HEK 293T/17 packaging cells using the Fugene 6 transfection reagent in DMEM High glucose complete media (DMEM High glucose supplemented with 10% FBS, Penicillin/Streptomycin, L-glutamine). 10 mM sodium butyrate was added to the transfected cells 3 hours post-transfection and left overnight. The following day this media was removed and the cells were washed briefly with 1X PBS before Opti-MEM media was added. Supernatants containing viral particles were collected every 12 hours between 36-72 hours after transfection, and these collections were pooled, filtered through 0.45 *μ*M SFCA or PES, and then concentrated by ultra-centrifugation. For infection, MEFs were incubated overnight with either mCherry-Lamin A or EYFP Lamin C viral supernatant and 4 *μ*g polybrene. Cells were allowed to expand in selection media containing 2 *μ*g/ml puromycin or 20 *μ*g/ml blasticidin respectively for 2-4 days followed by a second round of transduction with the other fluorescent tagged lamin viral supernatant followed by expansion in selection media containing both 2 *μ*g/ml puromycin and 20 *μ*g/ml blasticidin.

## Author Contributions

X.W., V.E.H, and K.L.R. designed the project, X.W., V.E.H, J.C.H., M.G., and K.L.R. executed the experiments, X.W. analyzed and interpreted DamID data, X.W. designed live cell imaging systems including the modified LAD-tracer and fluorescent tagged proteins, and X.W. designed all shRNA and rescue constructs. Chromosome Conformation Paints were designed in collaboration with Agilent Biotechnologies. K.L.R., X.W. and V.E.H. wrote the the manuscript with input from J.C.H. All authors edited and approved the final manuscript.

## Acknowledgements

We thank Rakel Tryggvadottir and Sinan Ramazanoglu for invaluable assistance with sequencing and David Vinson for help with western blots. We are grateful to Jevon Cutler and Teresa Romeo Luperchio for great discussions and critiques that helped move this work forward. We thank the JHU Provost’s Office for support through the Catalyst and Discovery Awards, which served to support this work. This work was additionally funded through NIH from grants R01GM132427 and R21AG050132. M.G. was funded from NIH Training Grant T32GM007445 and V.E.H. partly funded from NIH training grant T32GM07814.

## Links to movies

Movie 1: EYFP-Lamin C and LAD tracer during interphase https://drive.google.com/file/d/1MZ6PDA-G4Zf4iAg0CYpuITuxJi5N6j2L/view?usp=sharing

Movie 2: EYFP-Lamin C and LAD tracer during interphase 2 https://drive.google.com/file/d/1HtdgRuwWY9KVm6eUrZT_nhIPIl57nu_6/view?usp=sharing

Movie 3: EYFP-Lamin C and LAD tracer during mitosis https://drive.google.com/file/d/1Wj_moDN5v0jQfCh1xyhtMmYLy6Zeu-yZ/view?usp=sharing

Movie 4: EYFP-Lamin C and LAD tracer during mitosis https://drive.google.com/file/d/1Zc2LTYamZyR0lgAgQ1VUOvH4EySEywkK/view?usp=sharing

## Supplemental Figures

**Supplemental Figure 1:**
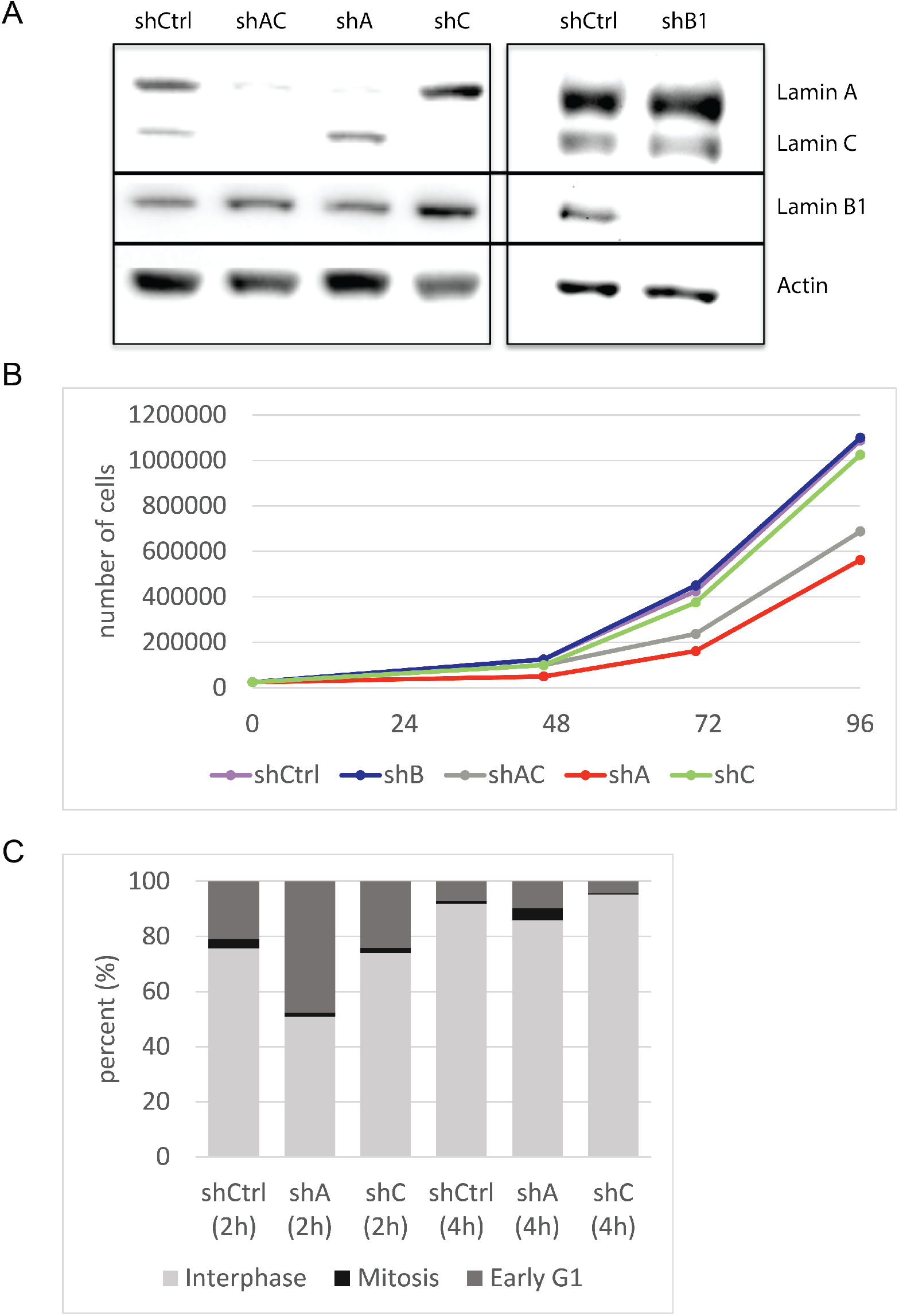
Analysis of shRNA knockdown cells. (A) Representative western blots showing specificity of the various knockdown constructs. (B) 25,000 cells were plated at 0 hours. Graph represents the number of cells on each plate after a given number of hours for each shRNA treatment indicated (shCtrl, shB1, shAC, shA, shC). (C) Graph indicates the percentage of synchronized cells in either interphase, mitosis or G1 as assessed from nuclear lamina morphology at either 2hours or 4hours after release into mitosis for each condition (shCtrl, shA or shC).

**Supplemental Figure 2:**
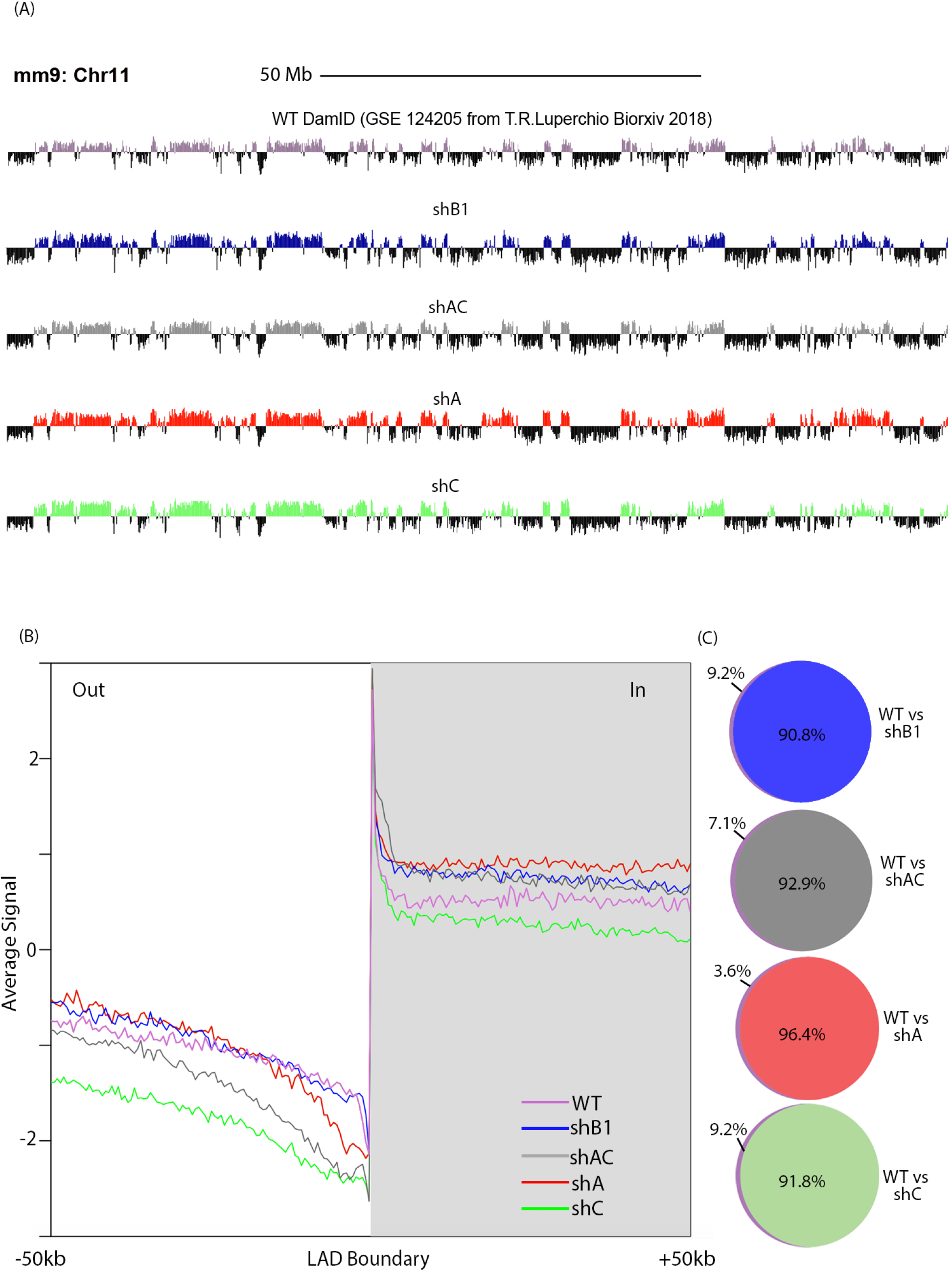
DamID analysis of shRNA treated cells. (A) Chromosome wide DamID traces (chromosome 11; chr 11) for wild-type (purple), lamin B1 depleted (blue), lamin A/C depleted (gray), lamin A depleted (red), and lamin C depleted (green) samples. Vertical axes are of log_2_ scale and traces above 0 (in indicated color), depict a higher than expected frequency of peripheral association. (B)Profiles of aligned LAD border regions (left and mirrored right border regions combined) are shown for lamin B1 interactions with chromatin (DNA). To align LAD borders, genome-wide positions of lamin B1 interactions were converted to coordinates relative to the nearest border. Gray area and positive coordinates, inside LADs; white area and negative coordinates, outside LADs. (C)Venn Diagram showing the proportion of wild type LAD domains preserved in the absence of lamin B1 or lamin A or C alone or both lamins A and C.

**Supplemental Figure 3a:**
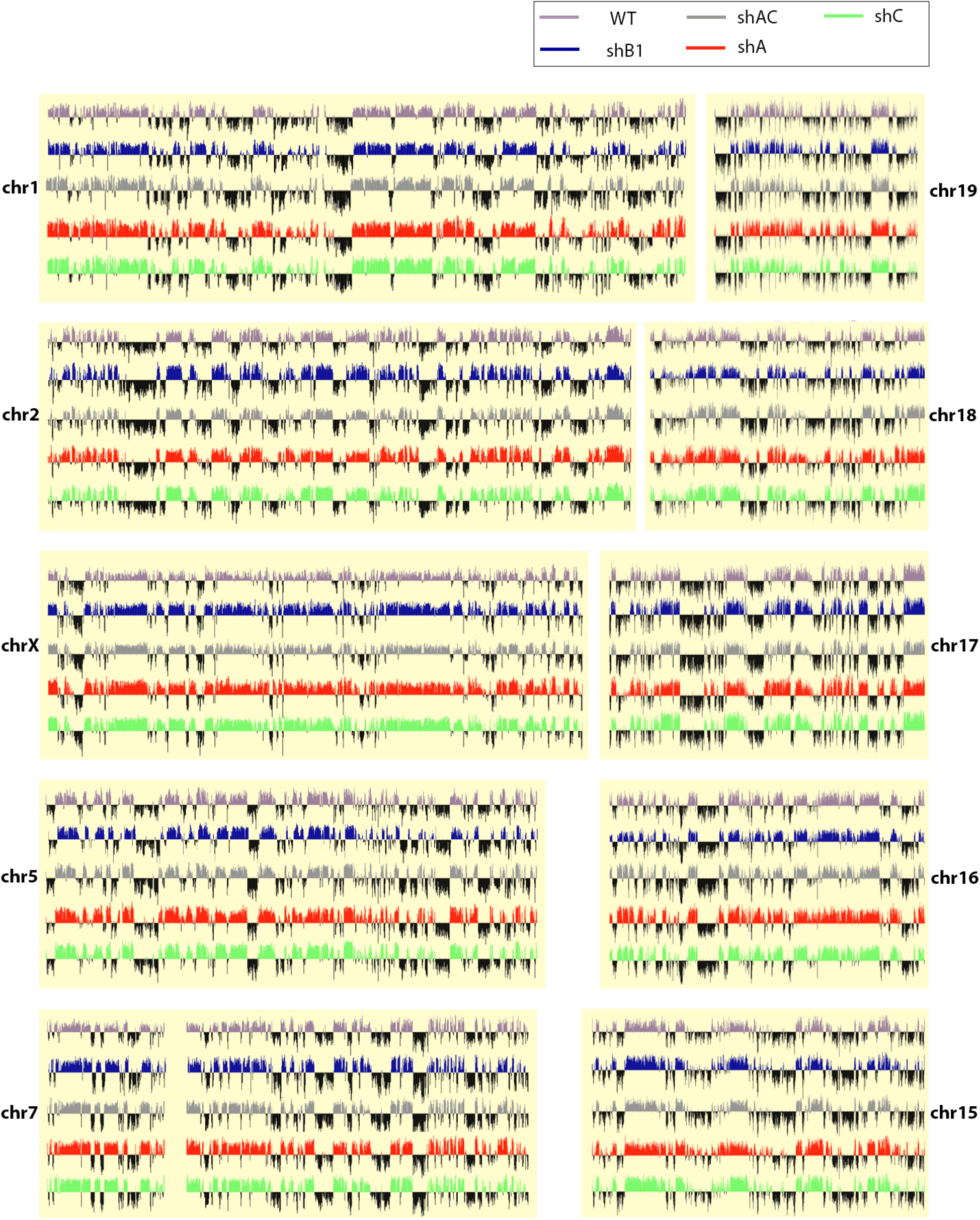
Genome-wide normalized DamID signal. Chromosome wide DamID traces for wild-type (purple), lamin B1 depleted (blue), lamin A/C depleted (gray), lamin A depleted (red), and lamin C depleted (green) samples. Vertical axes are of log_2_ scale and traces above 0 (in indicated color), depict a higher than expected frequency of peripheral association.

**Supplemental Figure 3b:**
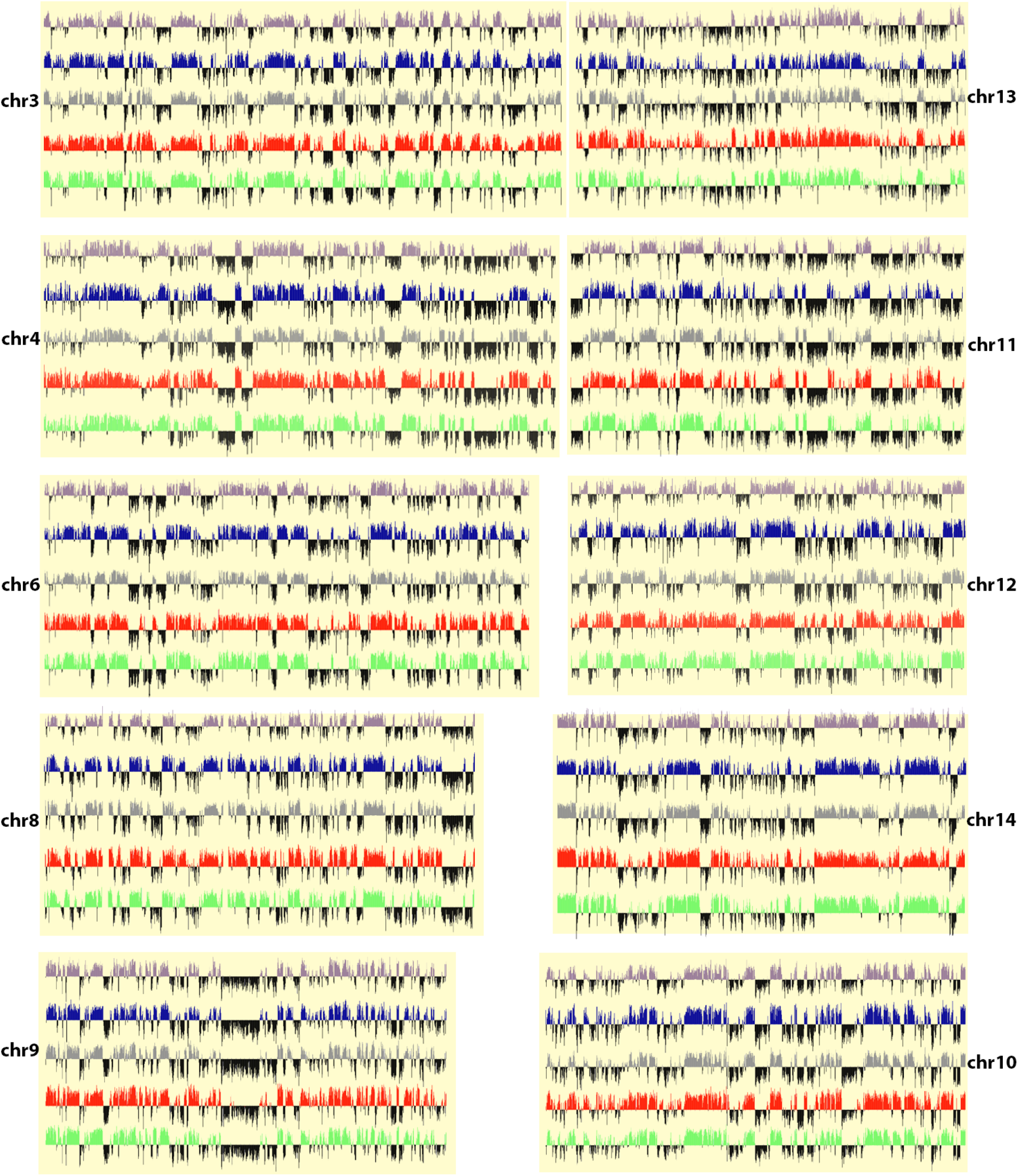
Genome-wide normalized DamID signal. Chromosome wide DamID traces for wild-type (purple), lamin B1 depleted (blue), lamin A/C depleted (gray), lamin A depleted (red), and lamin C depleted (green) samples. Vertical axes are of log_2_ scale and traces above 0 (in indicated color), depict a higher than expected frequency of peripheral association.

**Supplemental Figure 4:**
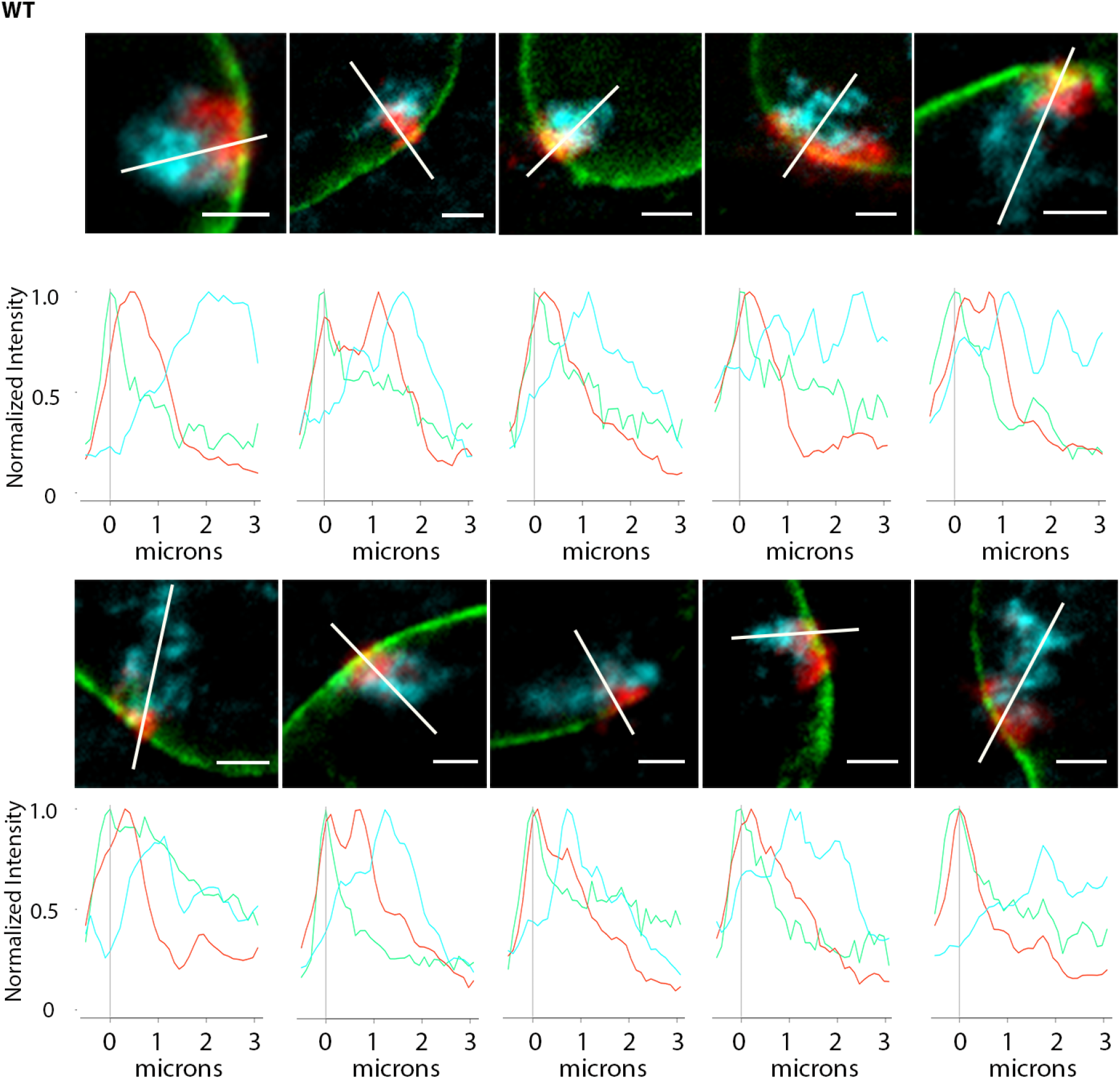
Chromosome conformation paints for chromosome 11 in WT (untreated) MEFs. Representative images of chromosome 11 conformation paints to chromosome nonLADs (cyan) LADs (red) lamin B1 (green). Normalized fluorescence intensity histogram plots for chromosome 11 territories in wild type MEFs, plotted from the nuclear lamina (0*μ*m) to 3*μ*m into the nucleus. The line each plot travels through is represented by a white line. Scale bar = 2*μ*m.

**Supplemental Figure 5:**
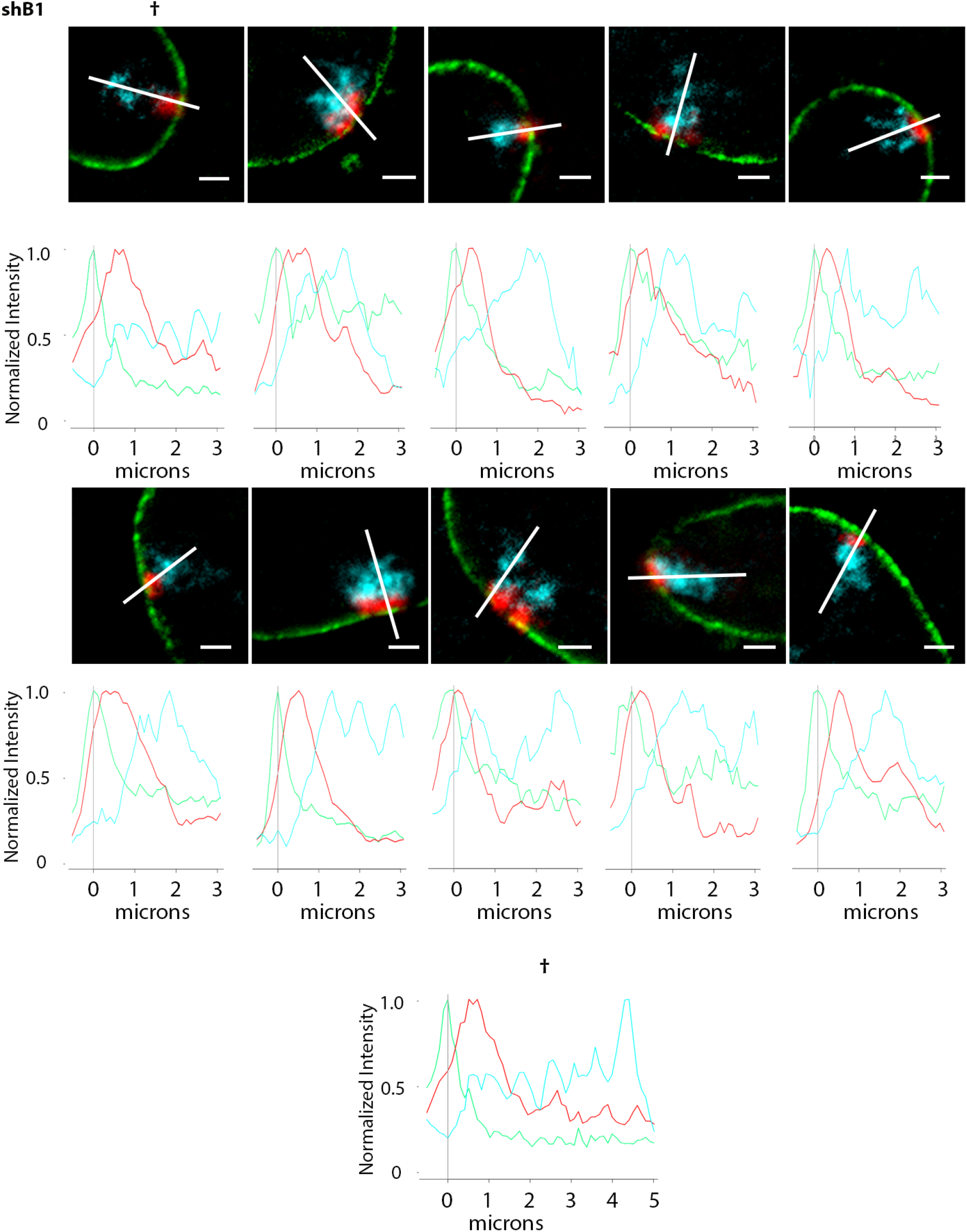
Lamin B1 depleted cells have normal LAD/nonLAD configuration. Representative images of chromosome 11 conformation paints to chromosome nonLADs (cyan) LADs (red) lamin A/C (green). Normalized fluorescence intensity histogram plots for chromosome 11 territories in shLB1 treated MEFs, plotted from the nuclear lamina (0*μ*m) to 3*μ*m into the nucleus. The line each plot travels through is represented by a white line. Scale bar = 2*μ*m.

**Supplemental Figure 6:**
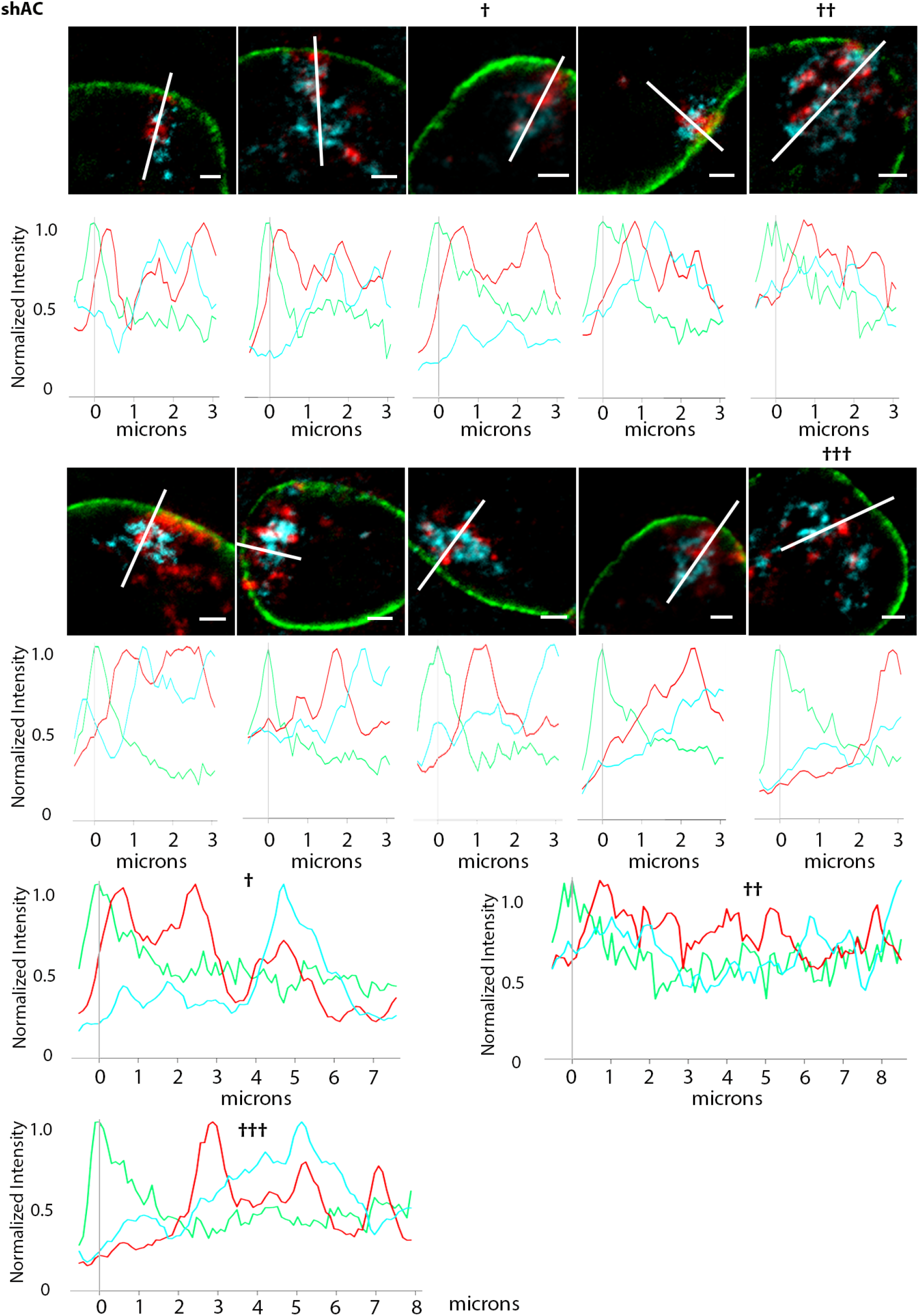
Lamin A/C depleted cells show wide-spread disruption of LAD and nonLAD organization. Representative images of chromosome 11 conformation paints to chromosome nonLADs (cyan) LADs (red) lamin B1 (green). Normalized fluorescence intensity histogram plots for chromosome 11 territories in shAC treated MEFs, plotted from the nuclear lamina (0*μ*m) to 3*μ*m into the nucleus. The territories marked with †, † †, or † † † were plotted beyond 3*μ*m to better capture the disposition of LADs/nonLADs. The line each plot travels through is represented by a white line. Scale bar = 2*μ*m.

**Supplemental Figure 7:**
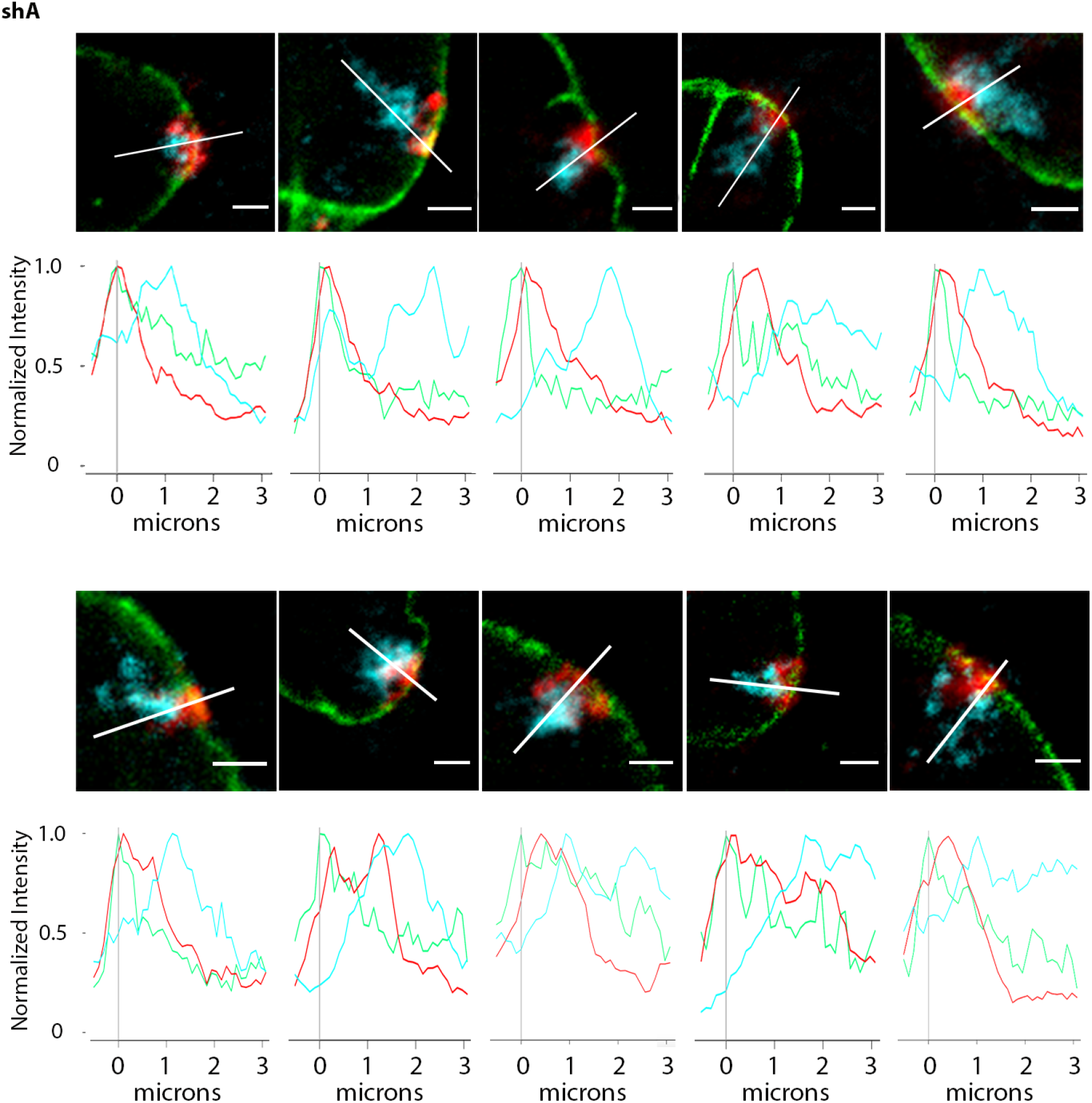
Lamin A (only) depleted cells have normal LAD/nonLAD configuration. Representative images of chromosome 11 conformation paints to chromosome nonLADs (cyan) LADs (red) lamin B1 (green). Normalized fluorescence intensity histogram plots for chromosome 11 territories in shA treated MEFs, plotted from the nuclear lamina (0*μ*m) to 3*μ*m into the nucleus. The line each plot travels through is represented by a white line. Scale bar = 2*μ*m.

**Supplemental Figure 8:**
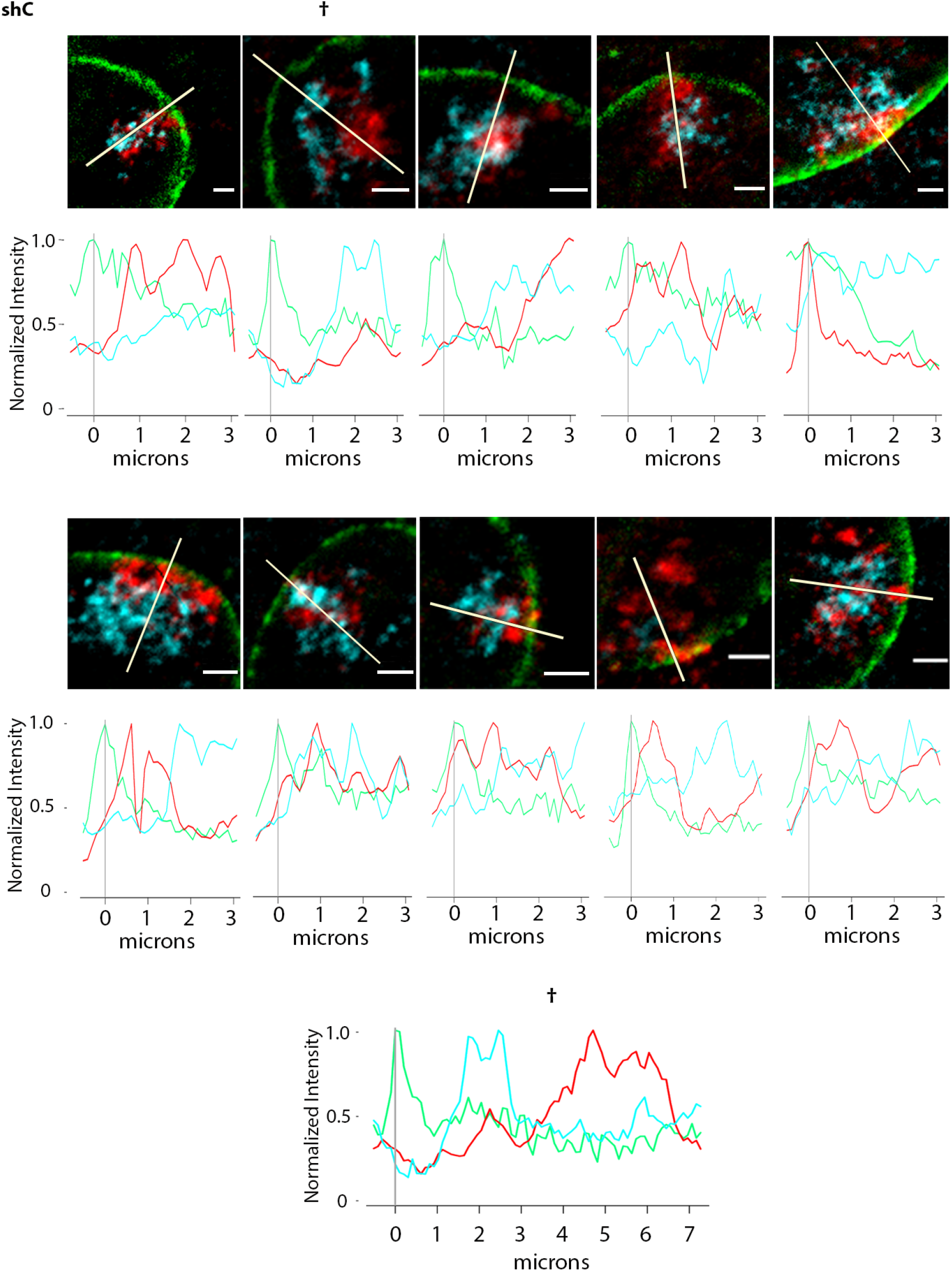
Lamin C (only) depleted cells show wide-spread disruption of LAD and nonLAD organization. Representative images of chromosome 11 conformation paints to chromosome nonLADs (cyan) LADs (red) lamin B1 (green). Normalized fluorescence intensity histogram plots for chromosome 11 territories in shC treated MEFs, plotted from the nuclear lamina (0*μ*m) to 3*μ*m into the nucleus. The territory marked with † was plotted beyond 3*μ*m to better display LAD and nonLAD signals. The line each plot travels through is represented by a white line. Scale bar = 2*μ*m.

**Supplemental Figure 9:**
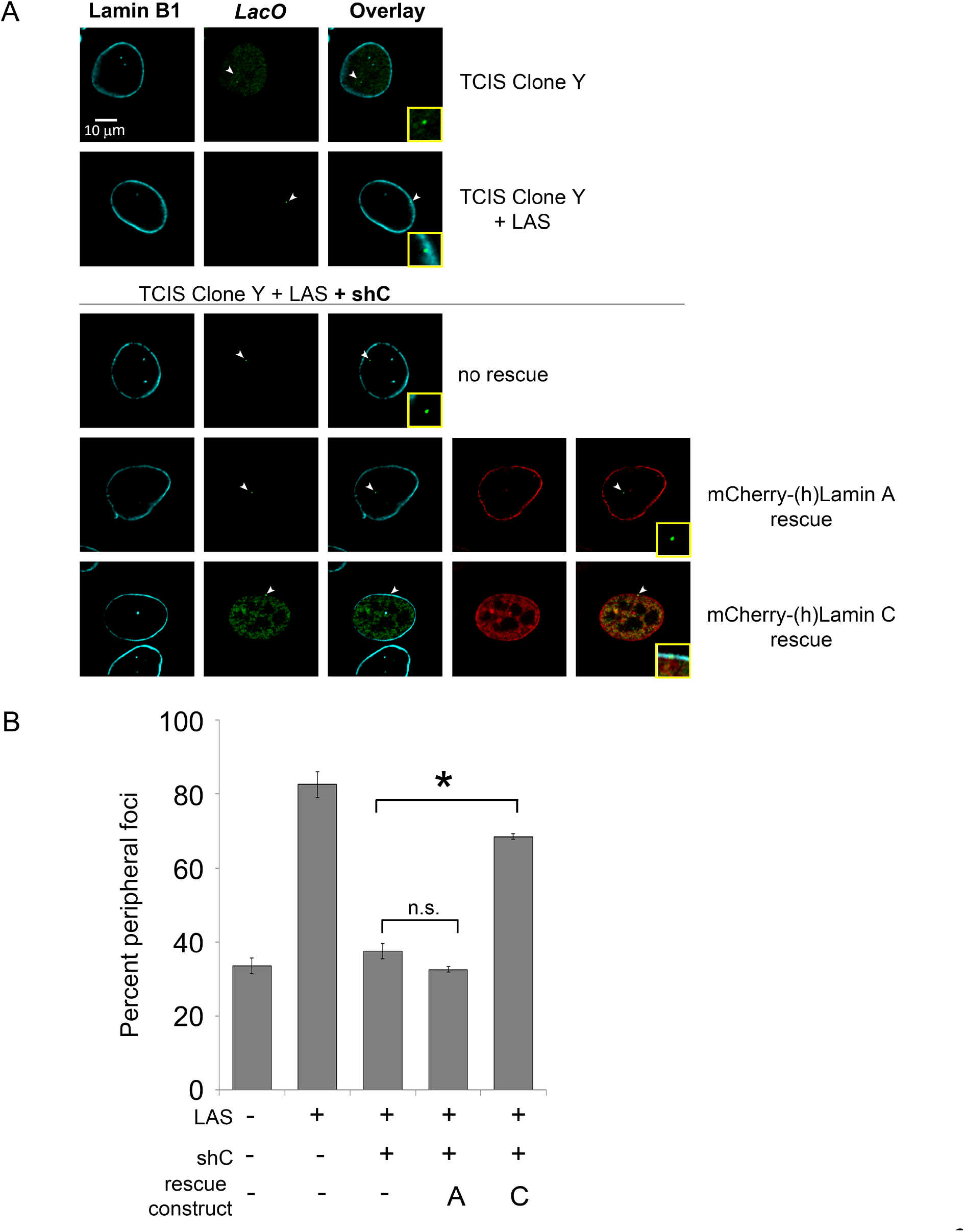
LAS localization can be rescued by expression of human lamin C. (A) Representative images showing the disposition of *lacO* arrays (arrowheads, green), lamin A or C respectively (red), and lamin B1 (cyan) in the TCIS clone Y. The inset shows 300× magnification. (B) Quantification of peripheral association was determined by the overlap of EGFP-LacI foci and lamin B1 (n ≥ 50). Error bars indicate SD. Asterisks for rescue experiment indicate significance of p ≤ 0.001, not significant is indicated by n.s.

**Supplemental Figure 10a:**
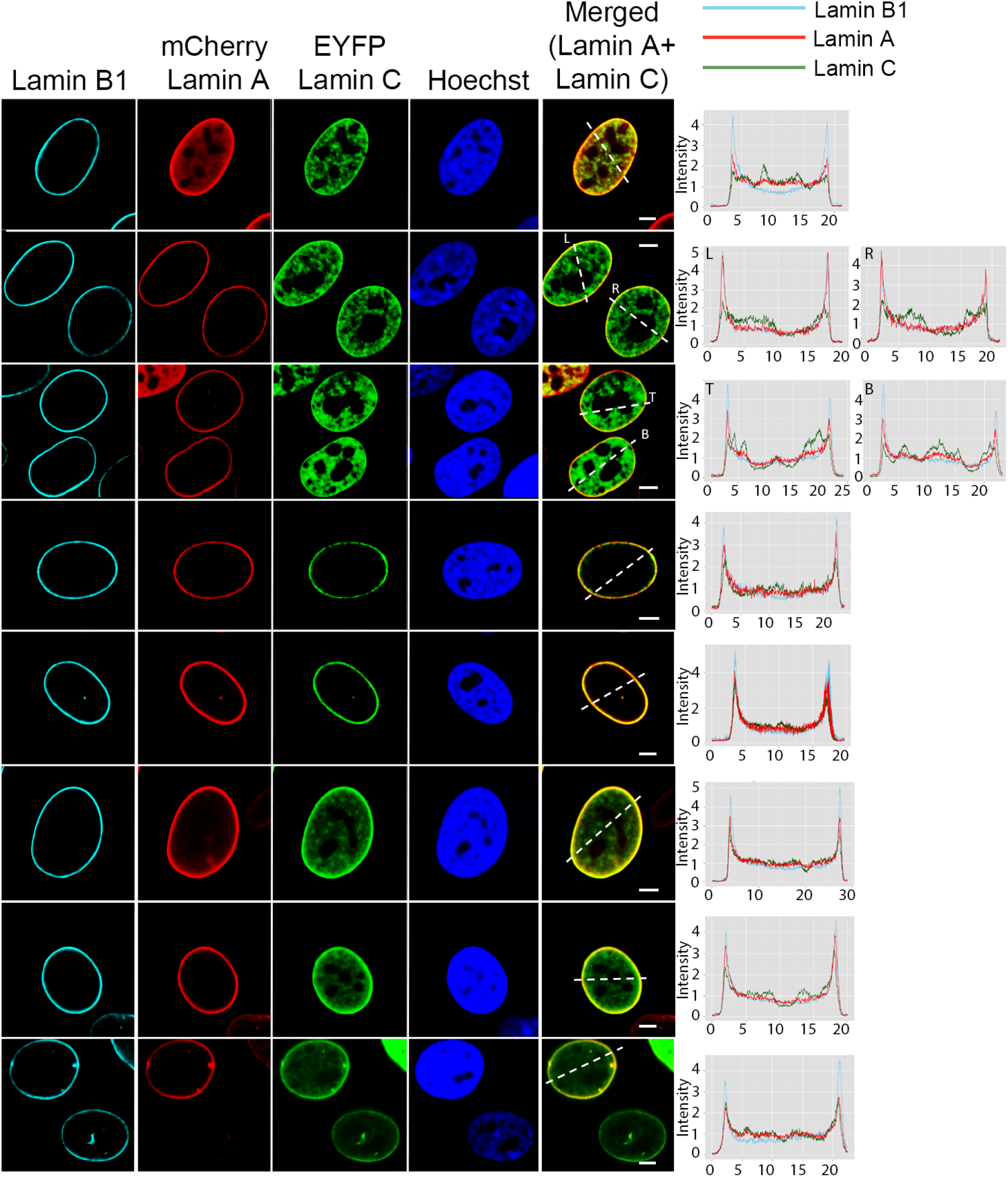
All 3 lamin isotypes localize to the periphery during interphase. Representative images of interphase nuclei anti-lamin B1 (cyan) mCherry-lamin A(red) EYFP-lamin C (green) Hoechst (blue). Fluorescence intensity histogram plots for lamin B1 (cyan), lamin A (red) and lamin C (green) along the indicated white dotted line in merge. Scale bar = 5*μ*m.

**Supplemental Figure 10b:**
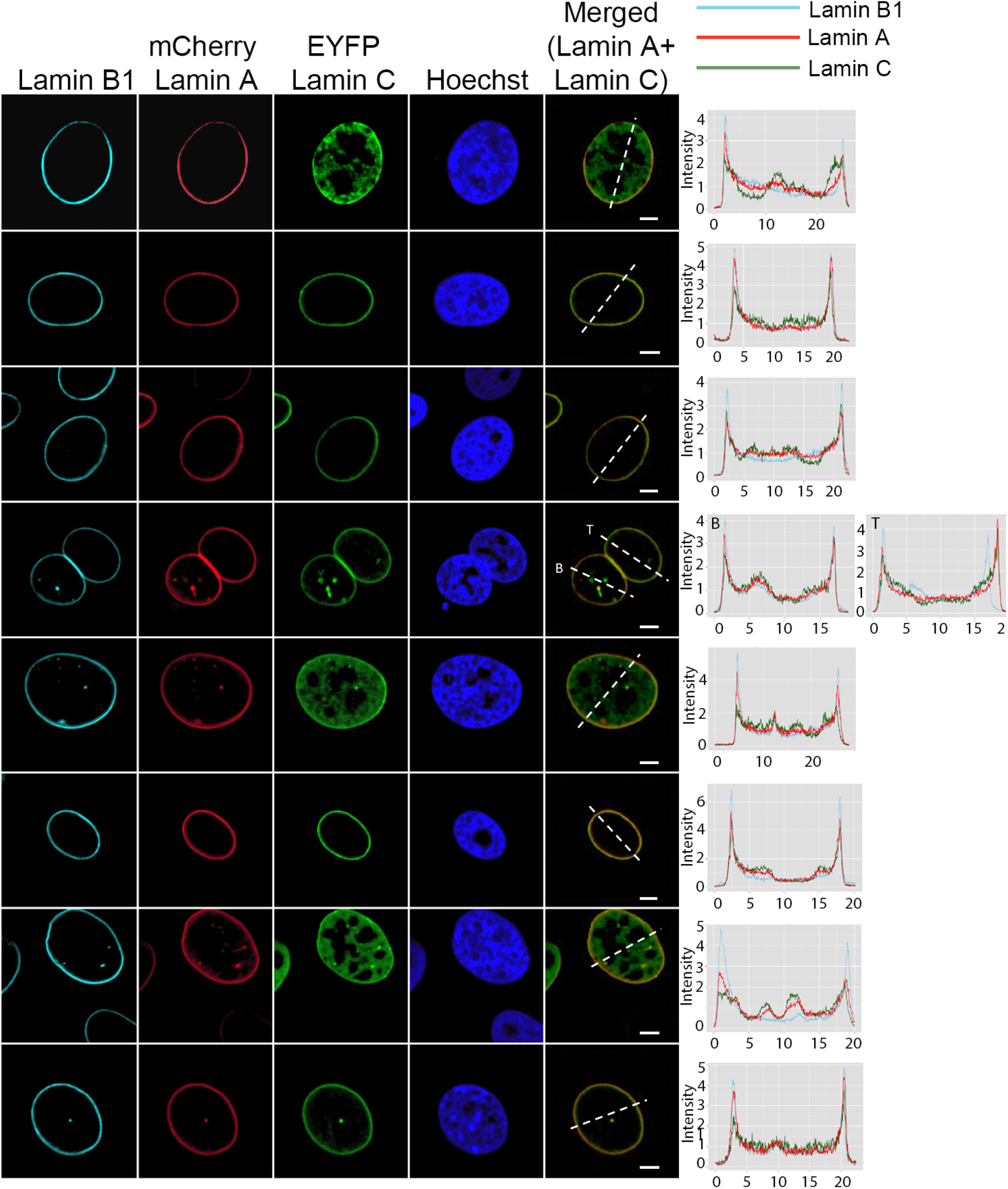
All 3 lamin isotypes localize to the periphery during interphase. Representative images of interphase nuclei anti-lamin B1 (cyan) mCherry-lamin A(red) EYFP-lamin C (green) Hoechst (blue). Fluorescence intensity histogram plots for lamin B1 (cyan), lamin A (red) and lamin C (green) along the indicated white dotted line in merge. Scale bar = 5*μ*m.

**Supplemental Figure 11a:**
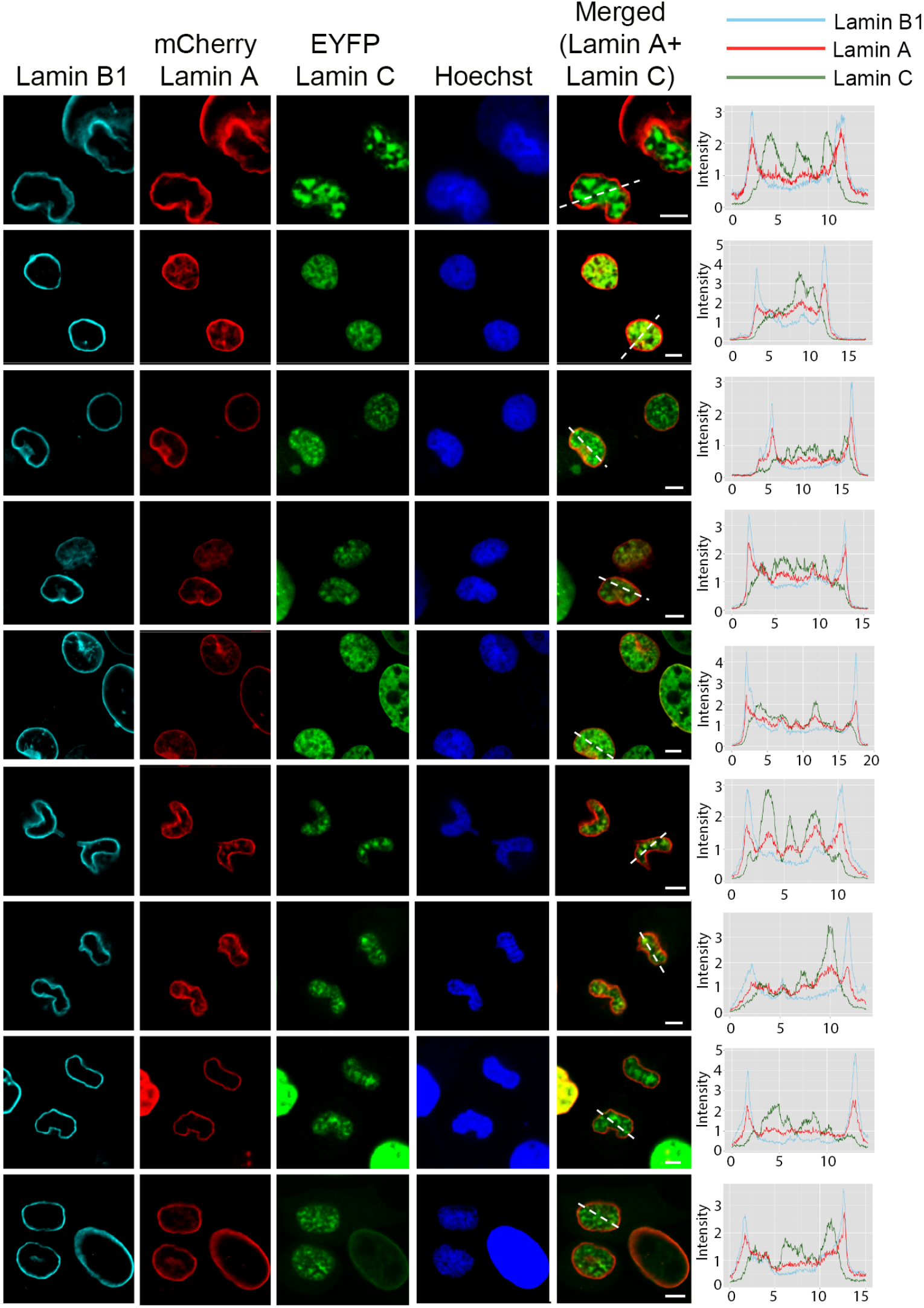
Differential localization of lamin B1, lamin A and lamin C at mitotic exit (telophase and early G1) Representative images of telophase and early G1 nuclei anti-lamin B1 (cyan) mCherry-lamin A (red) EYFP-lamin C (green) Hoechst (blue). Fluorescence intensity histogram plots for lamin B1 (cyan), lamin A (red) and lamin C green) along the indicated white dotted line in merge. Scale bar = 5*μ*m.

**Supplemental Figure 11b:**
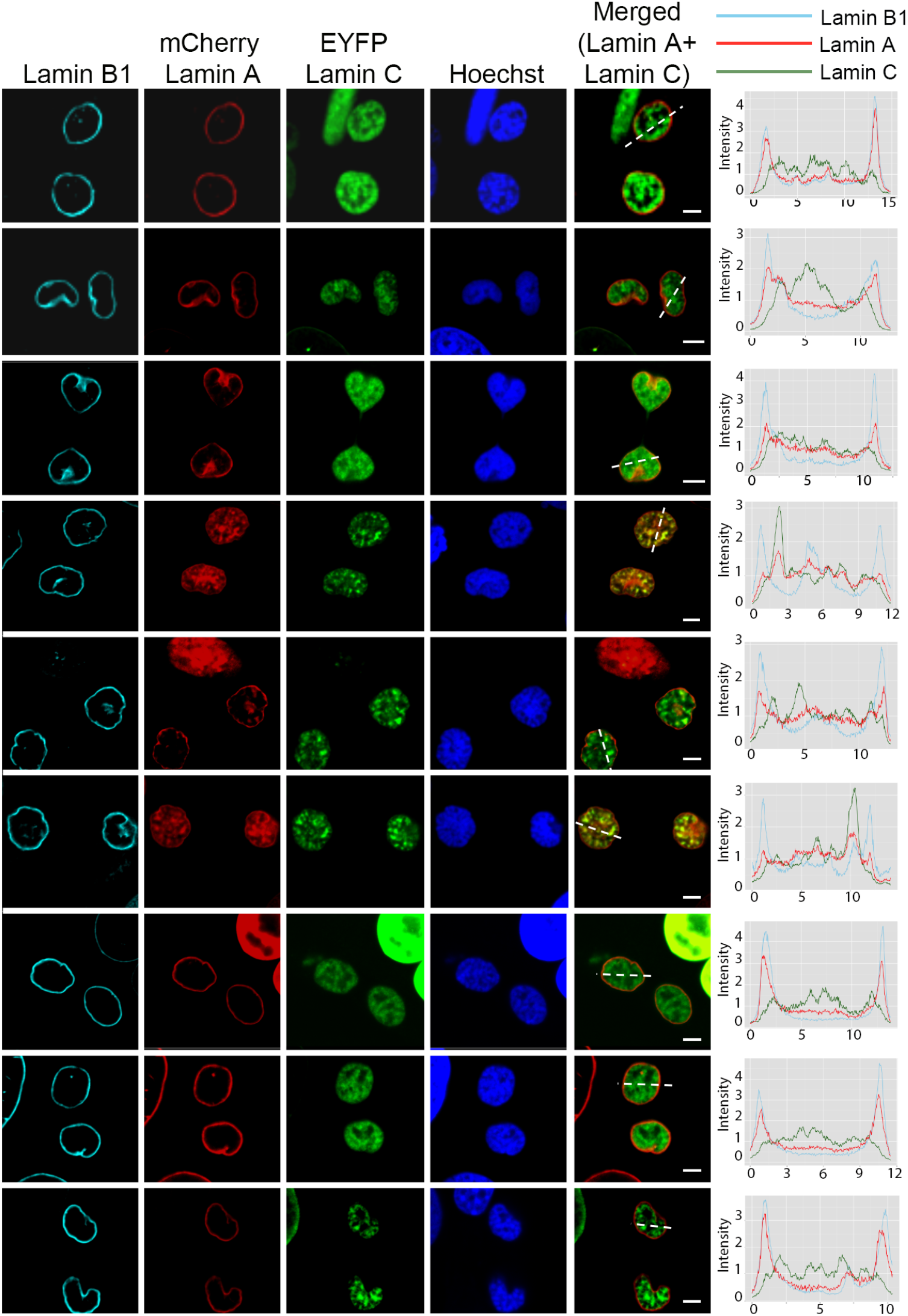
Differential localization of lamin B1, lamin A and lamin C at mitotic exit (telophase and early G1) Representative images of telophase and early G1 nuclei anti-lamin B1 (cyan) mCherry-lamin A(red) EYFP-lamin C (green) Hoechst (blue). Fluorescence intensity histogram plots for lamin B1 (cyan), lamin A (red) and lamin C (green) along the indicated white dotted line in merge. Scale bar = 5*μ*m.

**Supplemental Figure 12:**
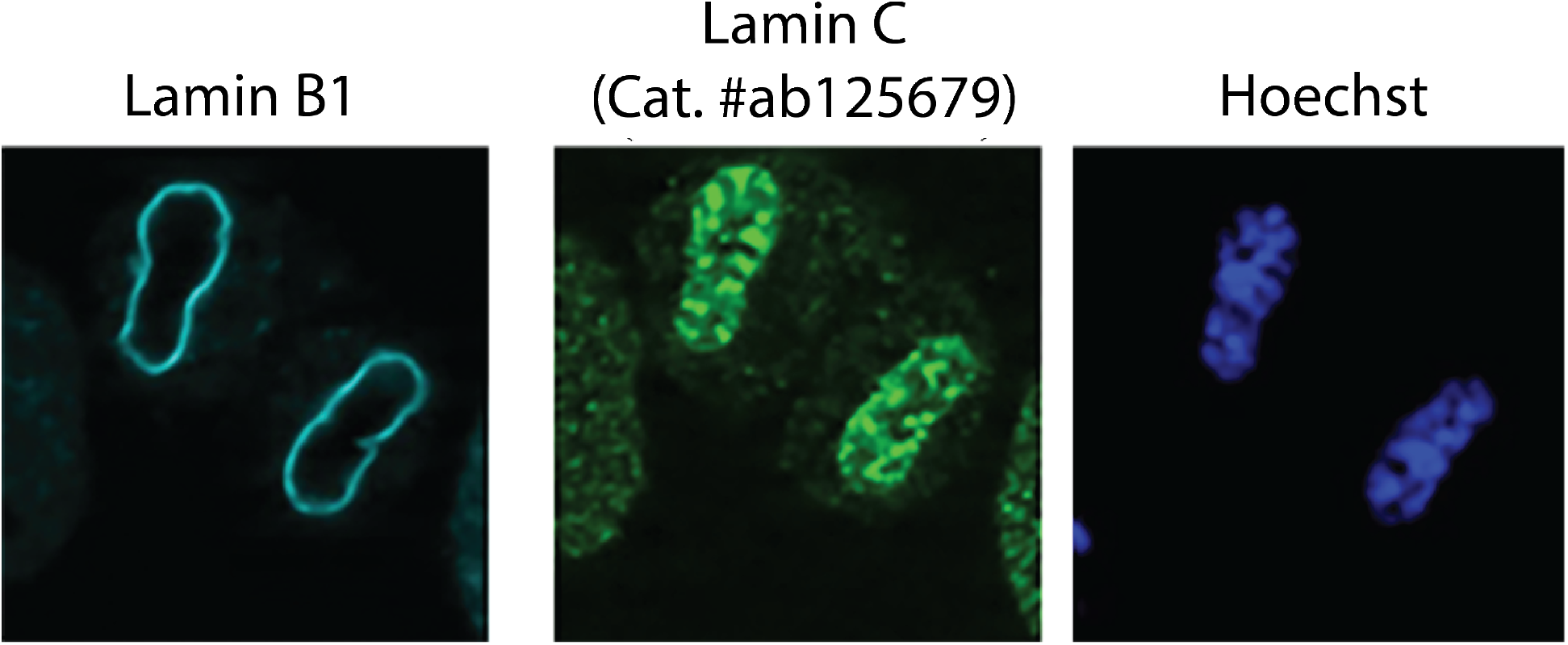
Antibody staining for lamin C shows the same localization as fluorescently tagged construct. Representative image of telophase/early G1 nucleus stained using antibody to lamin C (green, ab125679), antibody to lamin B1 (cyan) and Hoechst (blue).

**Supplemental Figure 13:**
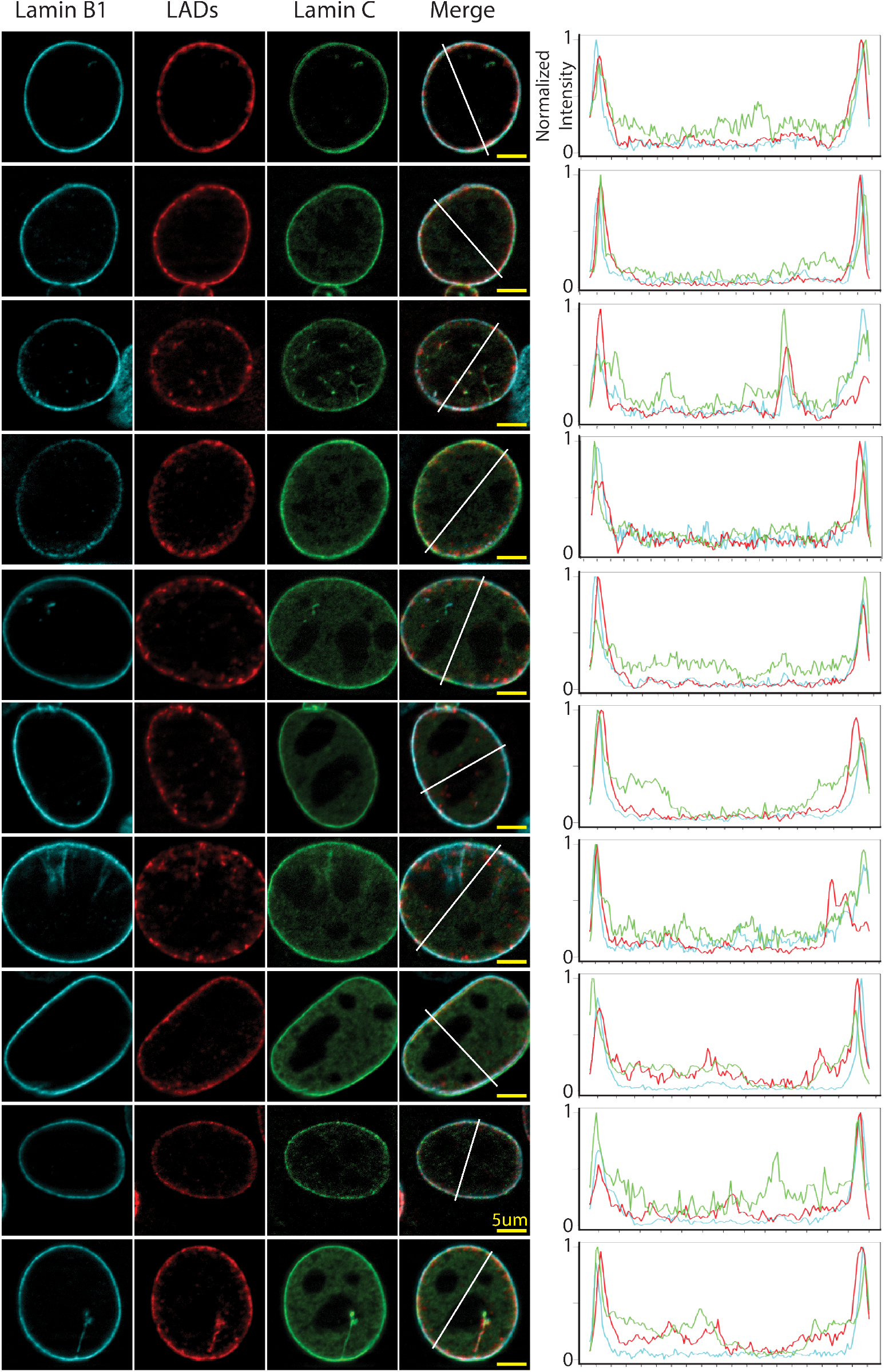
LADs and lamin C show peripheral localization during interphase. Representative images of interphase nuclei anti-lamin B1 (cyan) LAD-tracer (red) EYFP-lamin C (green). Normalized fluorescence intensity histogram plots for lamin B1 (cyan), LADs (red) and lamin C (green) along the indicated white line. Yellow scale bar = 5*μ*m.

**Supplemental Figure 14a:**
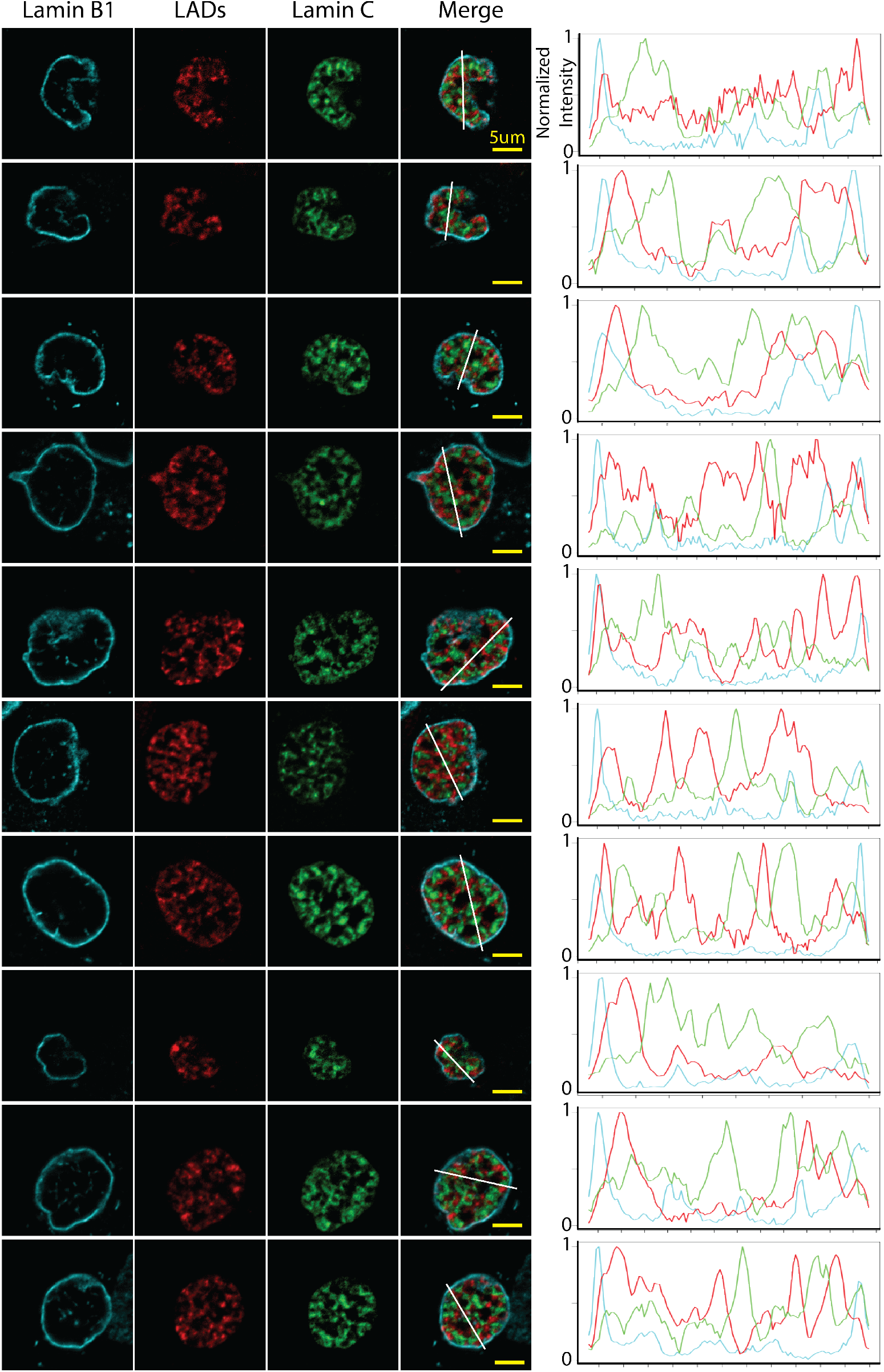
LADs and lamin C show nucleoplasmic localization during telophase/early G1 but do no co-localize. Representative images of telophase or early G1 nuclei anti-lamin B1 (cyan) LAD-tracer (red) EYFP-lamin C (green). Normalized fluorescence intensity histogram plots for lamin B1 (cyan), LADs (red) and lamin C (green) along the indicated white line. Yellow scale bar = 5*μ*m.

**Supplemental Figure 14b:**
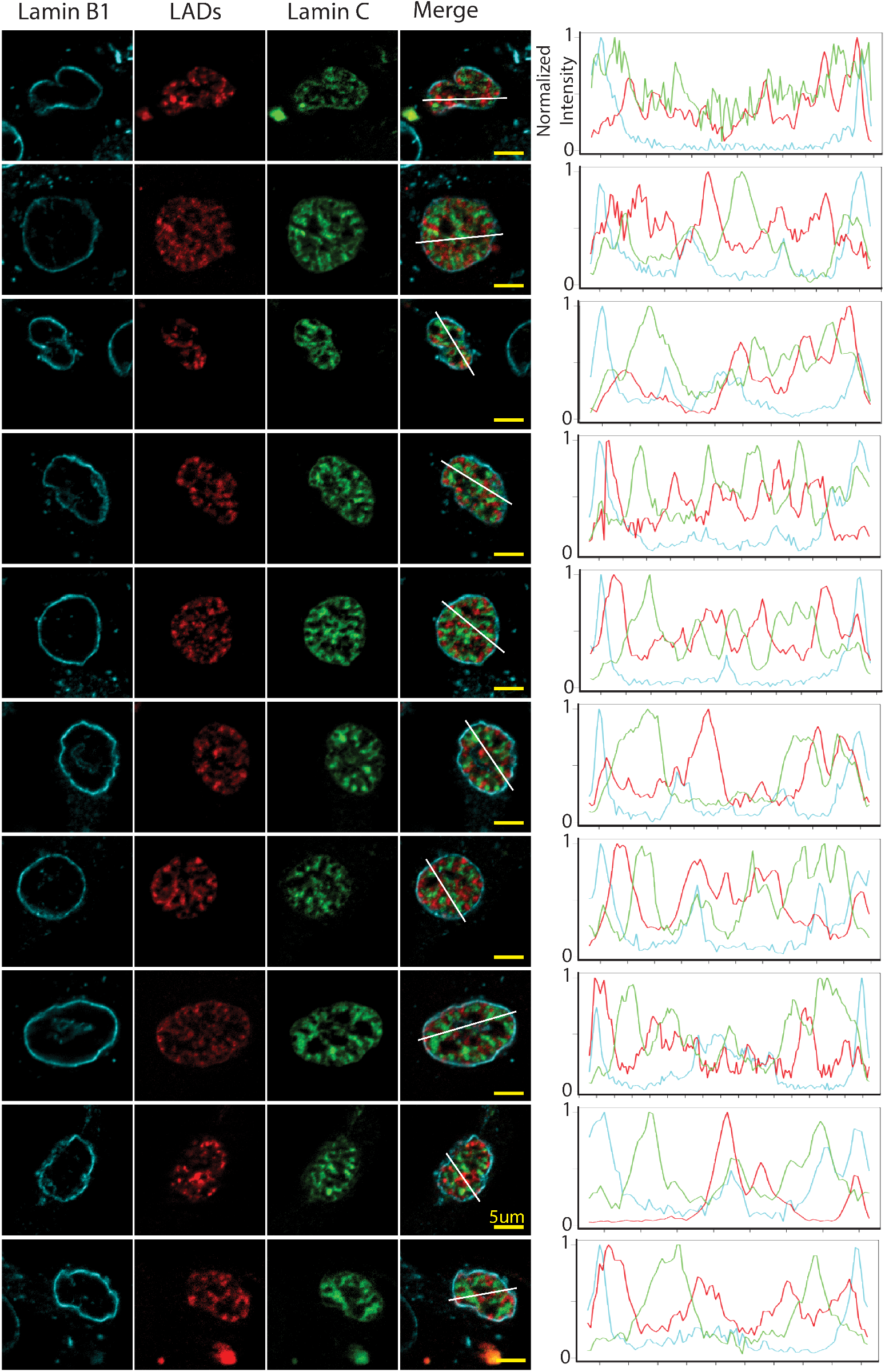
LADs and lamin C show nucleoplasmic localization during telophase/early G1 but do not co-localize. Representative images of telophase or early G1 nuclei anti-lamin B1 (cyan) LAD-tracer (red) EYFP-lamin C (green). Normalized fluorescence intensity histogram plots for lamin B1 (cyan), LADs(red) and lamin C (green) along the indicated white line. Yellow scale bar = 5*μ*m.

**Supplemental Figure 15:**
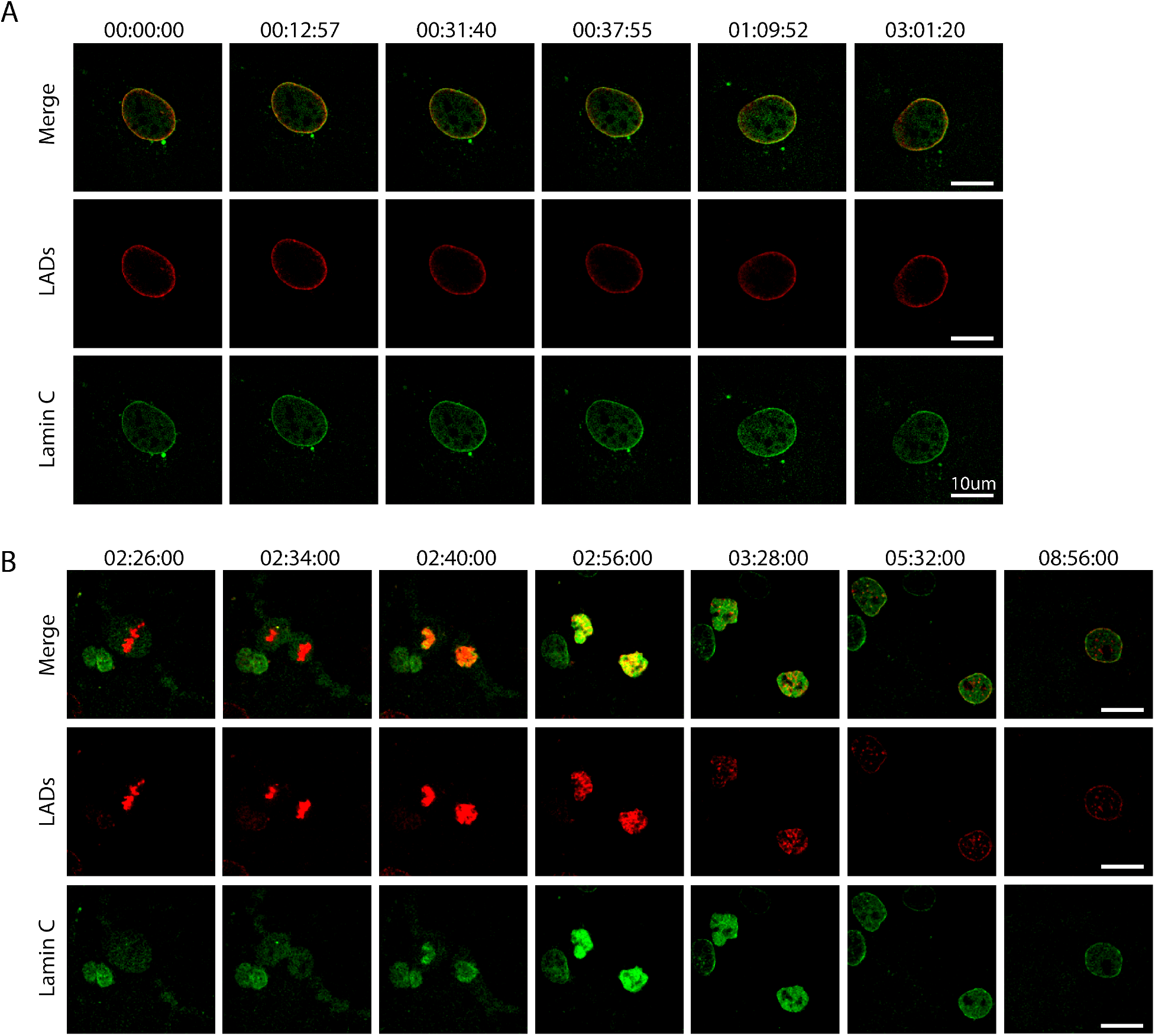
Lamin C and LADs resolve concurrently during G1. (A) Still images from time lapse movie 2 of LADs (red) EYFP-lamin C (green) during interphase shown over a similar time scale to movies 1 and 3. Scale bar is 20*μ*m. (B) Still images from time lapse movie 4 of LADs (red) EYFP-lamin C (green) during mitosis. Scale bar is 20*μ*m. Images were chosen to exemplify certain stages (metaphase, anaphase, telophase, early G1, partially resolved, fully resolved, mid G1). This movie extends further in time than other movies showing how LAD configuration continues to become tighter to the lamina well into G1 phase.

**Supplemental Figure 16a:**
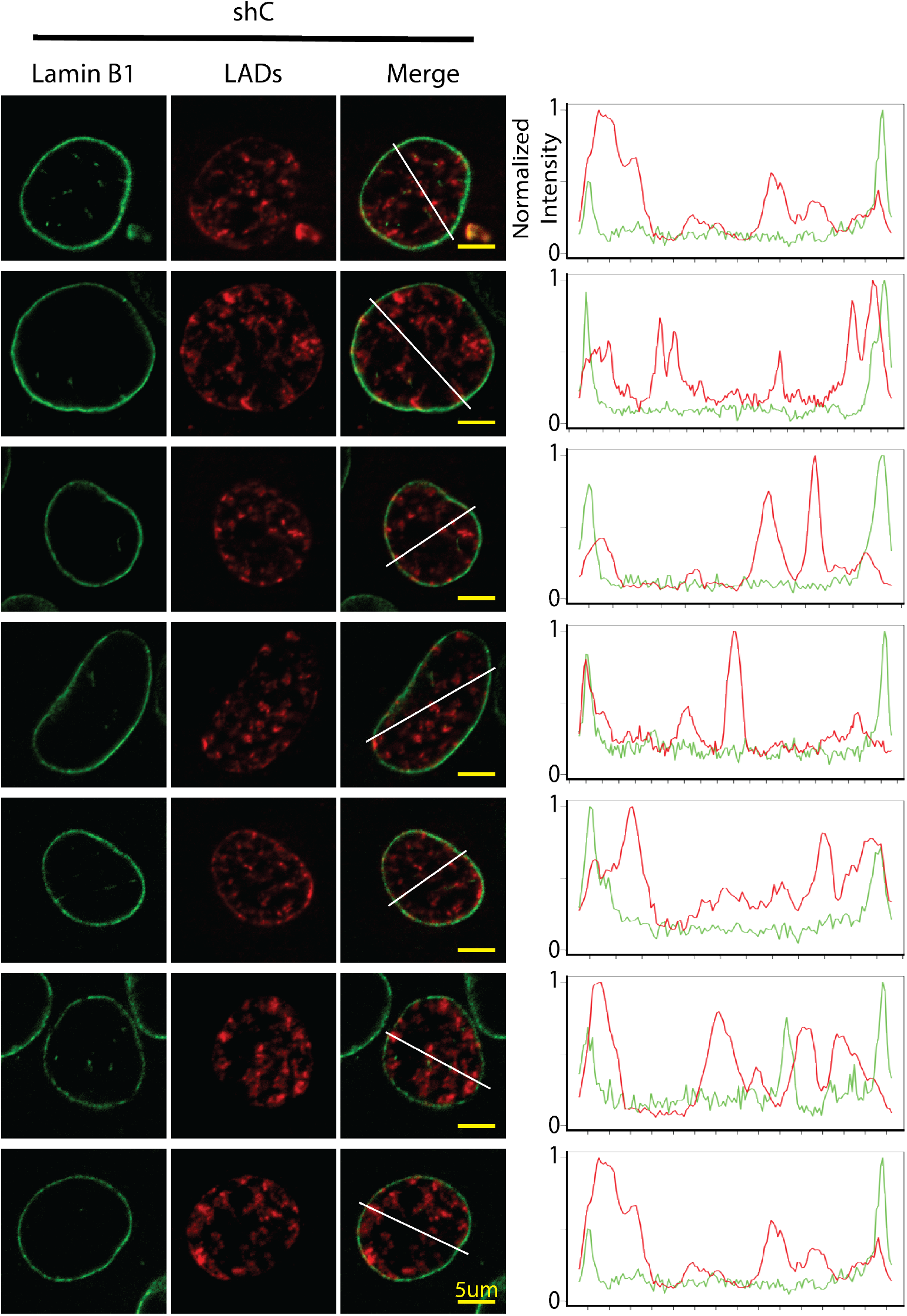
LADs form network like structures after cell division in the absence of lamin C. Representative images of cells harboring the LAD-tracer system that were transduced with a short hairpin RNA to lamin C. Lamin B1 (green) LADs (red). Normalized fluorescence intensity histogram plots for lamin B1 (green) and LAD-tracer (red) along the indicated white line. Yellow scale bar = 5*μ*m.

**Supplemental Figure 16b:**
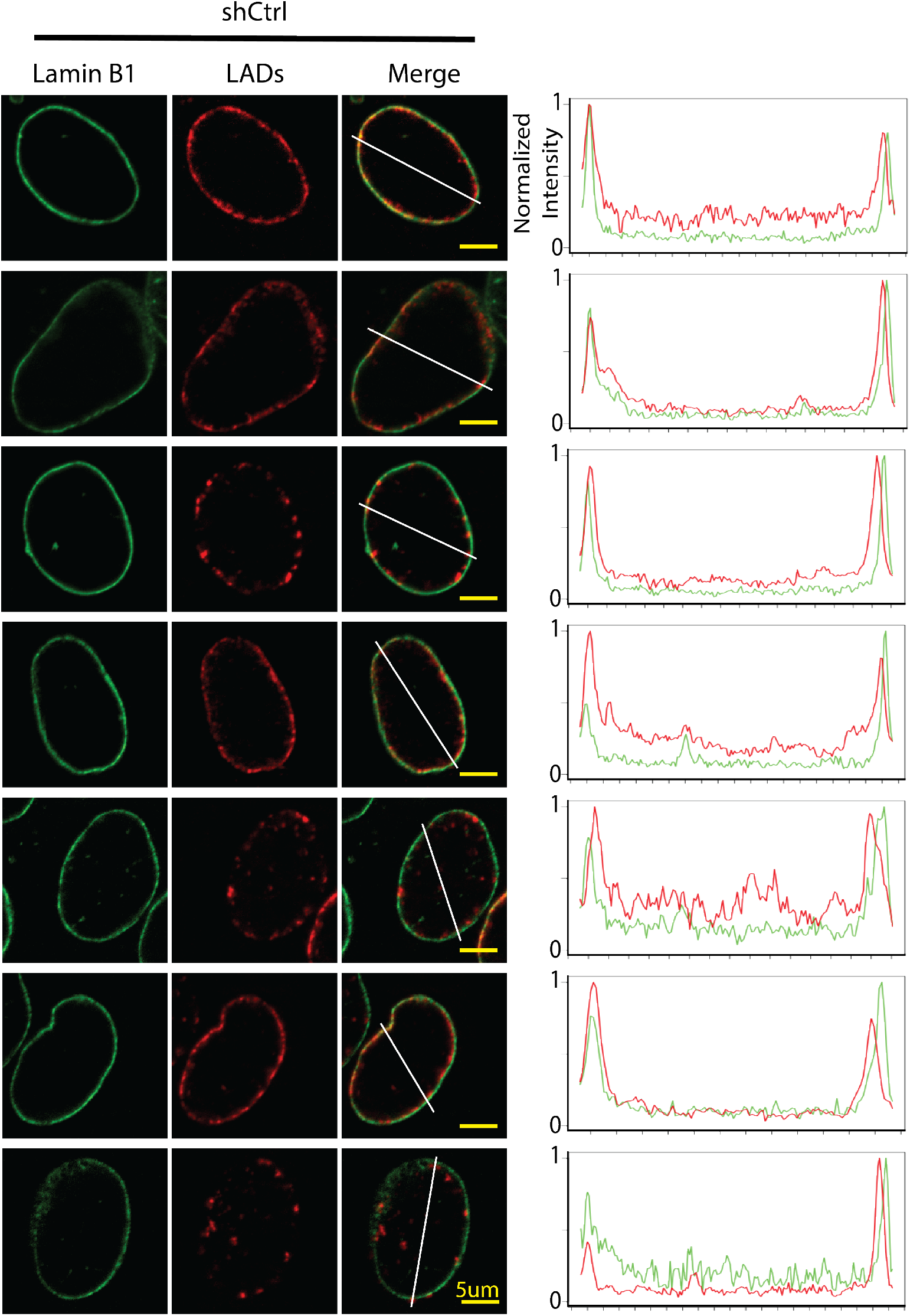
LADs reposition normally to the NE with a control shRNA. Representative images of cells harboring the LAD-tracer system that were transduced with a short hairpin RNA to a non-specific target. Lamin B1 (green) LADs (red). Normalized fluorescence intensity histogram plots for lamin B1 (green) and LAD-tracer (red) along the indicated white line. Yellow scale bar = 5*μ*m.

